# Constitutive AP2γ deficiency reduces postnatal hippocampal neurogenesis and induces behavioral deficits in juvenile mice that persist during adulthood

**DOI:** 10.1101/2021.06.07.447328

**Authors:** Eduardo Loureiro-Campos, Nuno Dinis Alves, António Mateus-Pinheiro, Patrícia Patrício, Carina Soares-Cunha, Joana Silva, Vanessa Morais Sardinha, Bárbara Mendes-Pinheiro, Tiago Silveira-Rosa, Ana João Rodrigues, João Filipe Oliveira, Nuno Sousa, Luísa Pinto

## Abstract

The transcription factor activating protein two gamma (AP2γ) is an important regulator of neurogenesis both during embryonic development as well as in the postnatal brain, but its role for neurophysiology and behavior at distinct postnatal periods is still unclear. In this work, we explored the neurogenic, behavioral, and functional impact of a constitutive AP2γ heterozygous deletion in mice from early postnatal development until adulthood. Constitutive AP2γ heterozygous deletion in mice caused a reduction of hippocampal transient amplifying progenitors (TAPs) in the postnatal brain, inducing significant impairments on hippocampal-dependent emotional- and cognitive-behavioral tasks including anxiety-like behavior and cognitive deficits, typically associated with an intact neurogenic activity. Moreover, AP2γ deficiency impairs dorsal hippocampus-to-prefrontal cortex functional connectivity.

We observed a progressive and cumulative impact of constitutive AP2γ deficiency on the hippocampal glutamatergic neurogenic process, as well as alterations on limbic-cortical connectivity, together with impairments on emotional and cognitive behaviors from juvenile to adult periods. Collectively, the results herein presented demonstrate the importance of AP2γ in the generation of glutamatergic neurons in the postnatal brain and its impact on behavioral performance.

## Introduction

New cells are continuously generated, differentiated into neurons, and integrated into the preexisting neural networks in restricted regions of the postnatal mice brain (Boldrini et al., 2018; Dennis et al., 2016; Kempermann et al., 2018; Moreno-Jiménez et al., 2019; Tobin et al., 2019). One of these so-called neurogenic niches is the subgranular zone (SGZ) of the hippocampal dentate gyrus (DG). Here, neural stem cells (NSC) give rise to mature neural cells including glutamatergic granular neurons in a fined and tuned process with many developmental steps sensitive to different regulatory influences (Kempermann et al., 2004; Mateus-Pinheiro et al., 2017; Tobin et al., 2019; Toda et al., 2019). Postnatal hippocampal glutamatergic neurogenesis exhibits a regulatory transcriptional sequence (Sox2→ Pax6→ Ngn2→ AP2γ→ Tbr2→ NeuroD→ Tbr1) that recapitulates the hallmarks of the embryonic glutamatergic neurogenic process in the cerebral cortex (Hochgerner et al., 2018; Mateus-Pinheiro et al., 2017; Nacher et al., 2005). Transcriptional factors as Pax6, Ngn2, Tbr2, NeuroD, and Tbr1 have several and distinct roles in proliferation, cell kinetics, fate specification, and axonal growth (Englund, 2005; Götz et al., 1998; Hevner, 2019; Hevner et al., 2006; Hochgerner et al., 2018).

Despite several efforts to understand the complex transcriptional network orchestration involved in the regulation of neurogenesis, both in early developmental stages and during adulthood, these are still to be fully understood (Bertrand et al., 2002; Brill et al., 2009; Englund, 2005; Hack et al., 2005; Hsieh, 2012; Mateus-Pinheiro et al., 2017; Waclaw et al., 2006). Recently, the transcription factor activating protein 2 gamma (AP2γ, also known as Tcfap2c or Tfa2c) was described to be an important regulator of glutamatergic neurogenesis in the adult hippocampus, being involved in the regulation of transient amplifying progenitors (TAPs) cells (Hochgerner et al., 2018; Mateus-Pinheiro et al., 2018, 2017). AP2γ belongs to the AP2 family of transcription factors that is highly involved in several systems and biological processes, such as cell proliferation, adhesion, developmental morphogenesis, tumor progression and cell fate determination (Eckert et al., 2005; Hilger-Eversheim et al., 2000; Thewes et al., 2010). In addition, AP2γ is functionally relevant during embryonic neocortical development, playing detrimental roles in early mammalian extraembryonic development and organogenesis (Pinto et al., 2009). AP2γ is critical for the specification of glutamatergic neocortical neurons and their progenitors, acting as a downstream target of Pax6 and being involved in the regulation of Tbr2 and NeuroD basal progenitors’ determinants (Mateus-Pinheiro et al., 2017; Pinto et al., 2009). Strikingly, in humans, defects in the AP2γ gene were reported in patients with severe pre- and post-natal growth retardation (Geneviève et al., 2005), and to be involved in the mammary, ovarian and testicular carcinogenesis (Hoei-Hansen et al., 2004; Li et al., 2002; Ødegaard et al., 2006).

AP2γ deletion during embryonic development results in a specific reduction of upper layer neurons in the occipital cerebral cortex, while its overexpression increases region- and time-specific generation of neurons from cortical layers II/III (Pinto et al., 2009). AP2γ expression persists in the adult hippocampus, particularly in a sub-population of TAPs acting as a positive regulator of the cell fate modulators, Tbr2 and NeuroD, and therefore as a promoter of proliferation and neuronal differentiation (Mateus-Pinheiro et al., 2017). Conditional and specific downregulation of AP2γ in the adult brain NSCs decreases the generation of new neurons in the hippocampal DG and disrupts the electrophysiological synchronization between the hippocampus and the medial prefrontal cortex (mPFC). Furthermore, mice with AP2γ conditional deletion exhibited behavioral impairments, particularly a deficient performance in cognitive-related tasks (Mateus-Pinheiro et al., 2018, 2017).

These studies reveal the crucial modulatory role of AP2γ during embryonic cerebral cortex development as well as its influence on glutamatergic neurogenesis and hippocampal-dependent behaviors during adulthood. Still, it is important to identify the longitudinal postnatal relevance of AP2γ to brain neurophysiology and behavior. Thus, in the present study, we explored the neurogenic, behavioral and functional impact of constitutive AP2γ heterozygous deletion in mice since early postnatal development until adulthood. We revealed a progressive impact of AP2γ deficiency on the hippocampal glutamatergic neurogenic process, alterations on limbic-cortical connectivity, accompanied by behavioral impairments on emotional and cognitive modalities from juvenile period to adulthood.

## Results

### Constitutive AP2γ deficiency decreases proliferation and neurogenesis in the postnatal DG, without affecting neuronal morphology

We sought to dissect the impact of constitutive and heterozygous deficiency of AP2γ in the modulation of postnatal neuronal plasticity in the hippocampus, including its effect in the hippocampal neurogenic niche and in the morphology of pre-existing DG granule neurons at different postnatal periods. In juvenile and adult mice, we assessed the expression of markers for different cell populations along the neurogenic process through western blot and immunofluorescence, and the morphology of granule neurons in the DG using the Golgi-Cox staining method (Figure 1A and B). We observed a significant decrease in the expression levels of AP2γ protein in the hippocampal DG of juvenile (Figure 1C) and adult (Figure 1D) AP2γ^+/−^ mice, with concomitant reduction of Pax6 and Tbr2 protein levels, but not Sox2, an upstream regulator of AP2γ (Mateus-Pinheiro et al., 2017).

**Figure 1:**
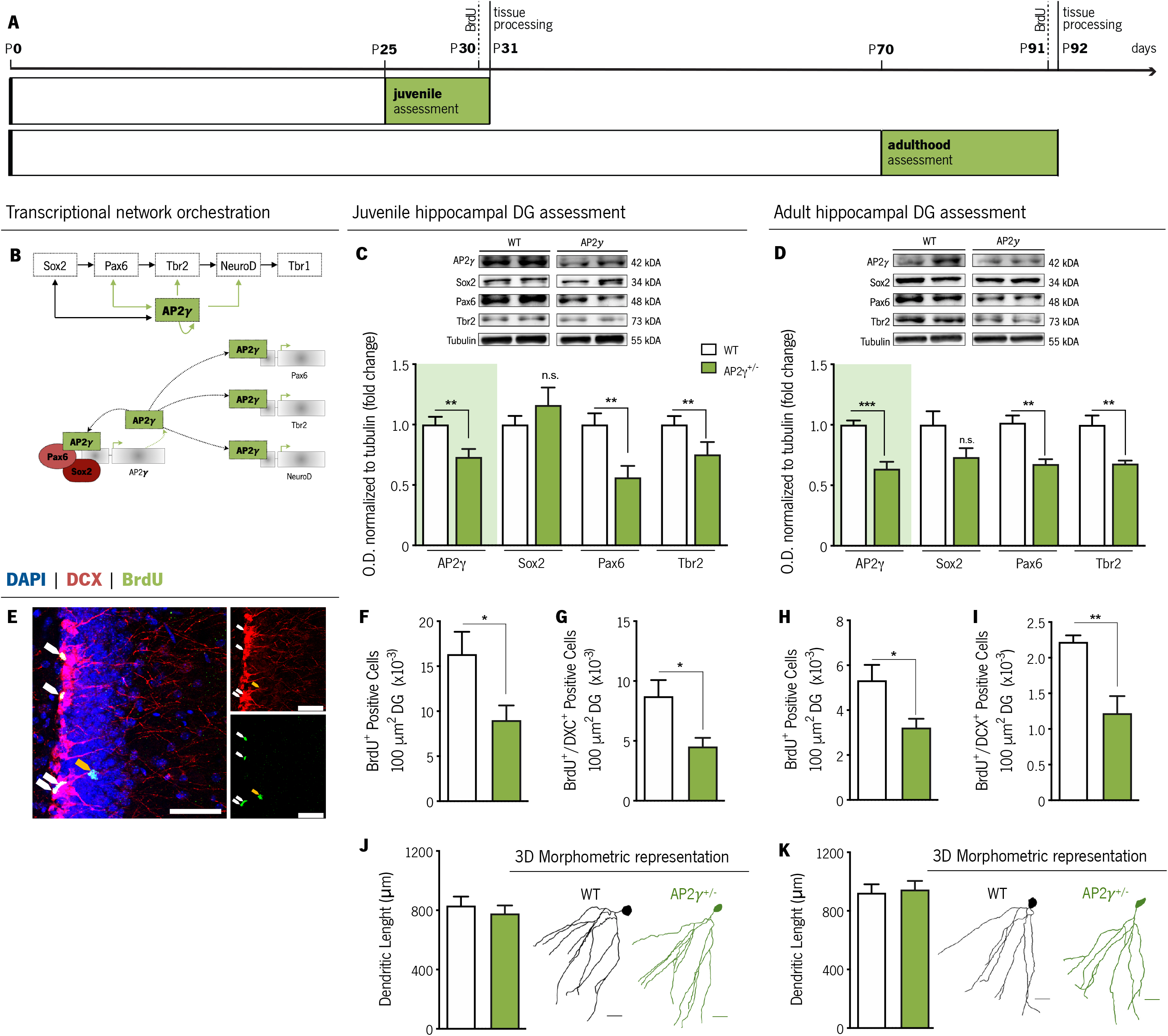
Constitutive and heterozygous AP2*y* deficiency reduces postnatal hippocampal neurogenesis both at juvenile and adult periods. (A) Experimental timeline. (B) Transcriptional network of hippocampal neurogenesis under modulatory role of AP2γ. Western-blot analysis of AP2γ, Sox2, Pax6, and Tbr2 in juvenile (C) and adult (D) dentate gyrus (DG) protein extracts. (E) Hippocampal DG coronal section immunostained for bromodeoxyuridine (BrdU) (green), doublecortin (DCX) (in red), and DAPI (in blue). BrdU/DCX double-positive cells are indicated by white arrows and solely BrdU-positive cell is identified with a yellow arrow. (F-I) Cell counts of BrdU-positive and BrdU/DCX double-positive cells in the hippocampal DG of juvenile and adult mice. (J and K) Dendritic length of three-dimensional (3D) neuronal reconstructed hippocampal granular neurons in juvenile (J) and adult (K) mice. Data presented as mean SEM. Sample size: Western-blot analysis: n_WT juvenile_ = 4; n_AP2γ^+/−^ juvenile_ = 4; n_WT adult_ = 4; n_AP2γ^+/−^ adult_ = 4; Immunostainings assays: n_WT juvenile_ = 6; n_AP2γ^+/−^ juvenile_ = 6; n_WT adult_ = 5; n_AP2γ^+/−^ adult_ = 5; 3D neuronal reconstruction: n_WT juvenile_ = 4; n_AP2γ^+/−^ juvenile_ = 4; n_WT adult_ = 4; n_AP2γ^+/−^ adult_ = 5. [Student’s t-test; \*\*\**p*<0.001, ** *p*< 0.01; * *p*< 0.05; Statistical summary in Supplementary table 1]. Scale bars represent 50 m. Abbreviations: WT, wild-type; AP2γ^+/−^, AP2γ heterozygous knockout mice; O.D., optical density.

Analysis of cell populations in the hippocampal neurogenic niche using BrdU and doublecortin (DCX) labelling (Figure 1E) revealed that the number of BrdU^+^ and BrdU^+^DCX^+^ cells is reduced in both juvenile and adult AP2γ^+/−^ mice, suggesting a decrease in the number of fast proliferating cells (transient amplifying progenitor cells (TAPs) and neuroblasts, respectively (Figure 1F-I). Of note, we observed a decrease in these cell populations with age in both WT and AP2γ^+/−^ mice in agreement with previous reports (Kase et al., 2020; Katsimpardi and Lledo, 2018) (BrdU^+^: Supplementary Figure 1A; BrdU^+^DCX^+^: Supplementary Figure 1B). In contrast, and despite an increased length (Supplementary Figure 1C) and neuronal arborization (Supplementary Figure 1D) of DG granular neurons with age, constitutive deficiency of AP2γ does not impact neither the dendritic length (Figure 1J-K), nor the neuronal complexity, in both postnatal periods (Supplementary Figure 1D).

These results highlight that AP2γ modulatory actions in the hippocampal neurogenic niche are similar and maintained in juvenile and adult mice. AP2γ transcription factor regulates NSCs proliferation and neuronal differentiation in the postnatal hippocampus by interacting with the different modulators involved in the transcriptional regulation of postnatal hippocampal neurogenesis.

### AP2γ^+/−^ mice display normal early postnatal development but anxiety-like behavior and cognitive impairments at juvenile period

In light of the negative impact of AP2γ heterozygous deficiency in the postnatal neurogenic process in the hippocampus, known as an important modulator of emotional and cognitive functions (Christian et al., 2014; Toda et al., 2019), we assessed early postnatal development and the behavioral performance of WT and AP2γ^+/−^ at juvenile and adult periods.

Evaluation of early postnatal development was performed through the assessment of somatic and neurobiological paraments during the first 21 postnatal days (Supplementary Figure 2). Despite a variation in the eye-opening day, responsiveness in sensory-motor functions, vestibular area-dependent tasks, and strength, as well somatic parameters were similar in WT and AP2γ^+/−^ mice. Furthermore, all analyzed parameters were within the previously described range (Guerra-Gomes et al., 2020; Heyser, 2003). These observations suggest that constitutive heterozygous deletion of AP2γ has no impact on early postnatal development.

In juvenile mice (between PND 25-31), we performed the open-field (OF) test to assess locomotor and anxiety-like behavior, tail suspension test (TST) and sucrose splash test (SST) to assess behavioral despair and anhedonic-like behavior, and the object recognition test (ORT) to assess memory (Figure 2A). In the OF, juvenile AP2γ^+/−^ mice exhibited a lower distance traveled in the anxiogenic center of the arena in comparison to WT animals (Figure 2B) suggesting an anxiety-like phenotype. Of note, WT and AP2γ^+/−^ mice display similar average velocities when performing the test, indicating no changes in locomotor activity (Supplementary Figure 3A). Assessment of behavioral despair and anhedonic-like behavior revealed no impact of constitutive AP2γ heterozygous deletion in these emotional domains, as no alterations were observed in the immobility time in the TST (Figure 2C) and grooming time in the SST (Figure 2D). Notably, despite no differences in the novel object location (Figure 2E and F), AP2γ^+/−^ mice displayed significant deficits in the novel object recognition, as denoted by a decreased preference to explore the novel object (Figure 2G).

**Figure 2:**
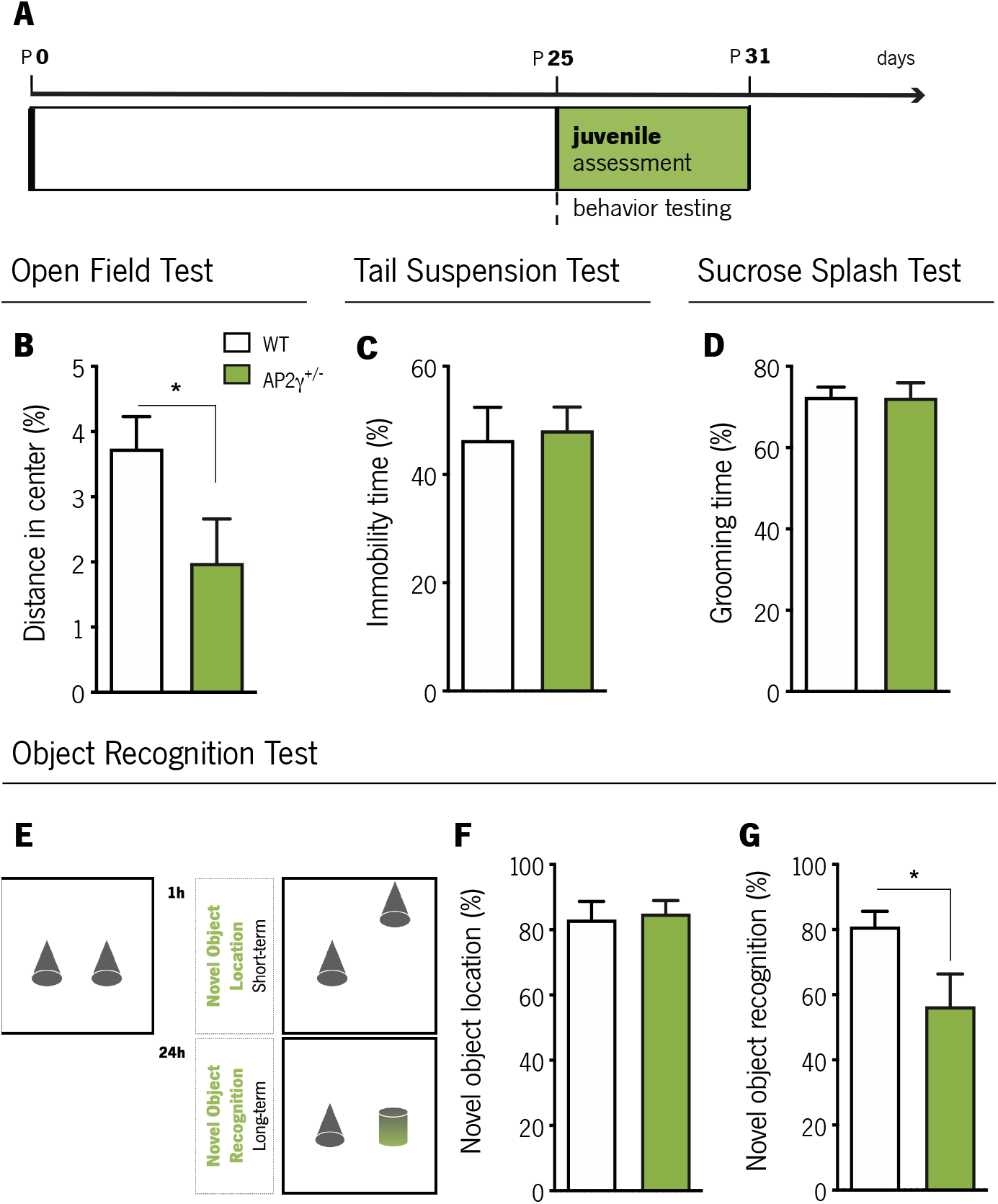
AP2γ deficiency increased anxiety-like behavior and promotes cognitive deficits in juvenile mice. (A) Timeline of behavioral assessment. Anxiety-like behavior was assessed through the open-field test (OF) (B), and depressive- and anhedonic-like behavior by the tail-suspension (TST) (C) and the sucrose splash (SST) (D) test. (E) To evaluate cognition, juvenile mice were subjected to the object recognition test, in particular to the novel object location (F) and the novel object recognition (G) tasks. Data presented as mean SEM. Sample size: OF: *n*_WT_ = 13; *n*_AP2γ^+/−^_ = 11; TST: *n*_WT_ = 16; *n*_AP2γ^+/−^_ = 13; ST: *n*_WT_ = 15; *n*_AP2γ^+/−^_ = 10; ORT: *n*_WT_ = 11; *n*_AP2γ^+/−^_ = 8. [Student’s t-test; * *p*< 0.05; Statistical summary in Supplementary table 1]. Abbreviations: WT, wild-type; AP2γ^+/−^, AP2γ heterozygous knockout mice.

These observations suggest that despite no evident impact on early postnatal development, constitutive AP2γ heterozygous deficiency leads to memory impairments and an anxious-like phenotype at juvenile period.

### AP2γ^+/−^ adult mice exhibit significant impairments in emotional and cognitive behavioral modalities

At adulthood, behavioral assessment included OF test and elevated plus-maze (EPM) to evaluate anxiety-like behavior, forced swimming test (FST) and TST to examine behavioral despair, and ORT, contextual fear conditioning (CFC) and Morris water maze (MWM) to evaluate cognitive performance (Figure 3A). In the OF test, AP2γ^+/−^ mice showed a trend towards a decrease in the distance traveled in the anxiogenic center of the arena (*p* = 0.05, Figure 3B), with no changes in locomotor activity as denoted by similar average velocity assessment (Supplementary Figure 4A). Moreover, in the EPM test, AP2γ^+/−^ mice spent significantly less time in the open-arms than WT mice (Figure 3C). AP2γ^+/−^ mice do not show any changes in behavioral despair since immobility times in FST (Figure 3D) and TST (Figure 3E) are identical to WT mice. These observations suggest that anxiety-like behavior promoted by constitutive AP2γ deficiency tend to persist in adult mice, with no alteration in behavioral despair.

**Figure 3:**
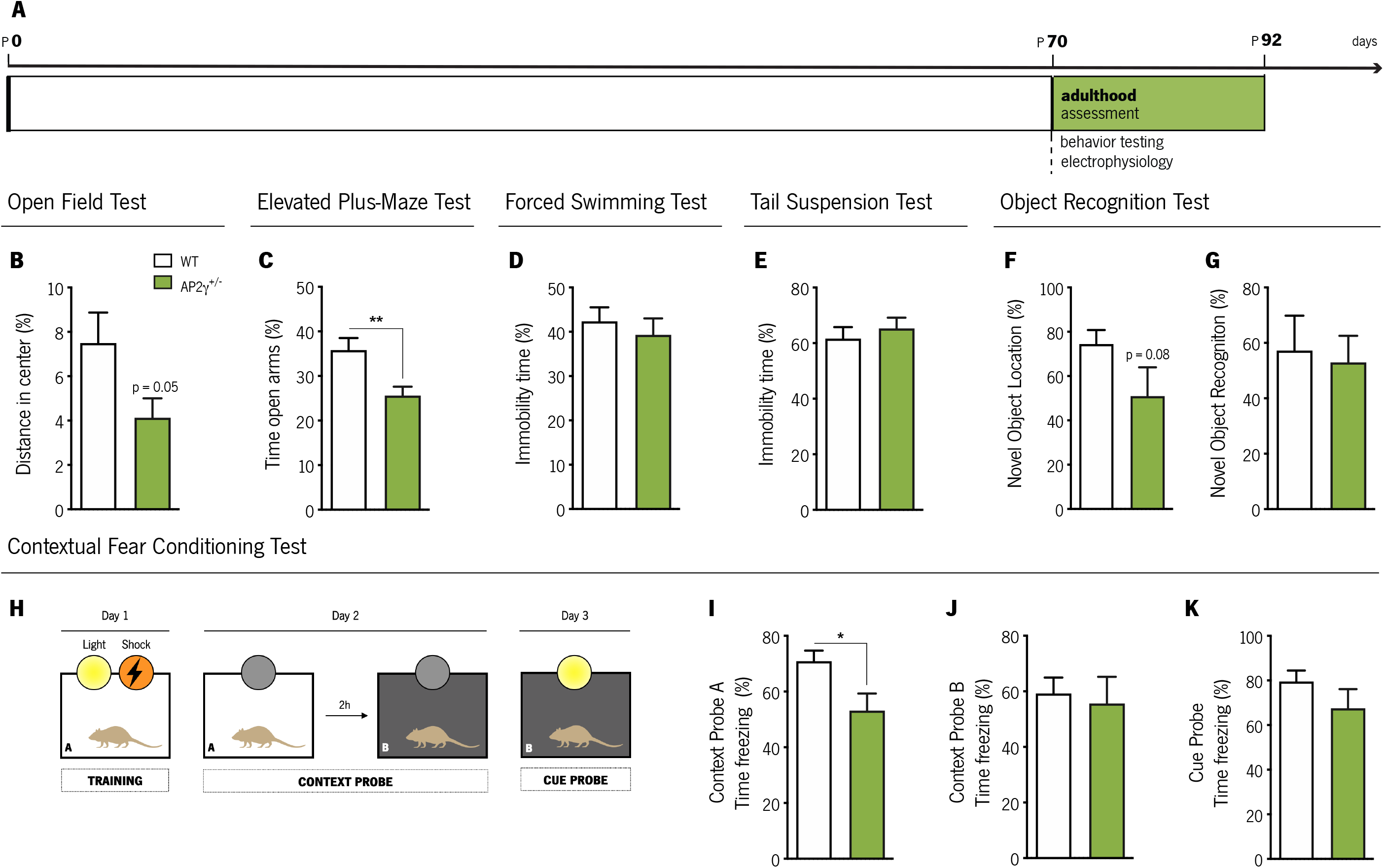
Behavioral assessment of adult mice. (A) Experimental timeline. (B and C) Anxiety-like behavior was assessed through the open-field test (OF) (B) and the elevated plus-maze (EPM) (C), while depressive-like behavior was evaluated through the forced-swimming test (FST) (D) and the tail-suspension (TST) (E) test. Object recognition test (ORT) (F and G) and (H-K) contextual fear conditioning (CFC) were performed to assess cognitive performance. Data presented as mean ± SEM. Sample size: OF, EPM and FST: *n*_WT_ = 12; *n*_AP2γ^+/−^_ = 14; TST: *n*_WT_ = 6; *n*_AP2γ^+/−^_ = 6; ORT: *n*_WT_ = 12; *n*_AP2γ^+/−^_ = 9; CFC: *n*_WT_ = 7; *n*_AP2γ^+/−^_ = 6. [Student’s t-test; ** *p*< 0.01; * *p*< 0.05; Statistical summary in Supplementary table 1]. Abbreviations: WT, wild-type; AP2γ^+/−^, AP2γ heterozygous knockout mice.

Cognitive performance assessed by ORT revealed a trend towards a decrease in the preference to explore the displaced object of AP2γ^+/−^ when compared to WT mice (*p* = 0.08, Figure 3F). However, no alterations were observed in preference towards the novel object (Figure 3G). In the CFC, a behavior test described to be sensitive to changes in adult hippocampal neurogenesis (Gu et al., 2012), mice were subjected to two distinct context tests, aimed to test hippocampal-dependent memory, and a cue probe to assess the integrity of extrahippocampal memory circuits (Figure 3H) (Gu et al., 2012; Mateus-Pinheiro et al., 2017). In the context A, AP2γ^+/−^ mice exhibited reduced freezing behavior when exposed to a familiar context (Figure 3I). No alterations in the freezing behavior were observed neither in the context B (Figure 3J) nor in the cue probe (Figure 3K). These observations suggest that AP2γ^+/−^ mice exhibit deficits in contextual hippocampal-related memory, and an intact associative non-hippocampal-dependent memory when compared to WT littermates. Furthermore, experimental groups were also subjected to the MWM test for evaluation of spatial memory (Figure 4). In the reference memory task, that relies on hippocampal function integrity (Cerqueira et al., 2007), AP2γ^+/−^ and WT mice exhibit similar performance to reach the hidden platform along the training days (Figure 4A and B). When the platform was changed to the opposite quadrant to assess behavior flexibility, which relies not only in the hippocampal formation but also in prefrontal cortical areas (Hamilton and Brigman, 2015), adult AP2γ^+/−^ mice spent less time in the new quadrant than WT animals (Figure 4C) suggesting that constitutive AP2γ deficiency leads to impaired behavioral flexibility. Detailed analysis of the strategies adopted to reach the escape platform (Antunes et al., 2020; Garthe et al., 2009; Garthe and Kempermann, 2013; Mateus-Pinheiro et al., 2017; Ruediger et al., 2012) revealed that AP2γ^+/−^ mice delayed the switch from non-hippocampal dependent ("Block 1”) to hippocampal-dependent (“Block 2”) strategies (Figure 4D-H), suggesting an impairment of hippocampal function. No differences were found in the working memory task (Supplementary Figure 4B and C).

**Figure 4:**
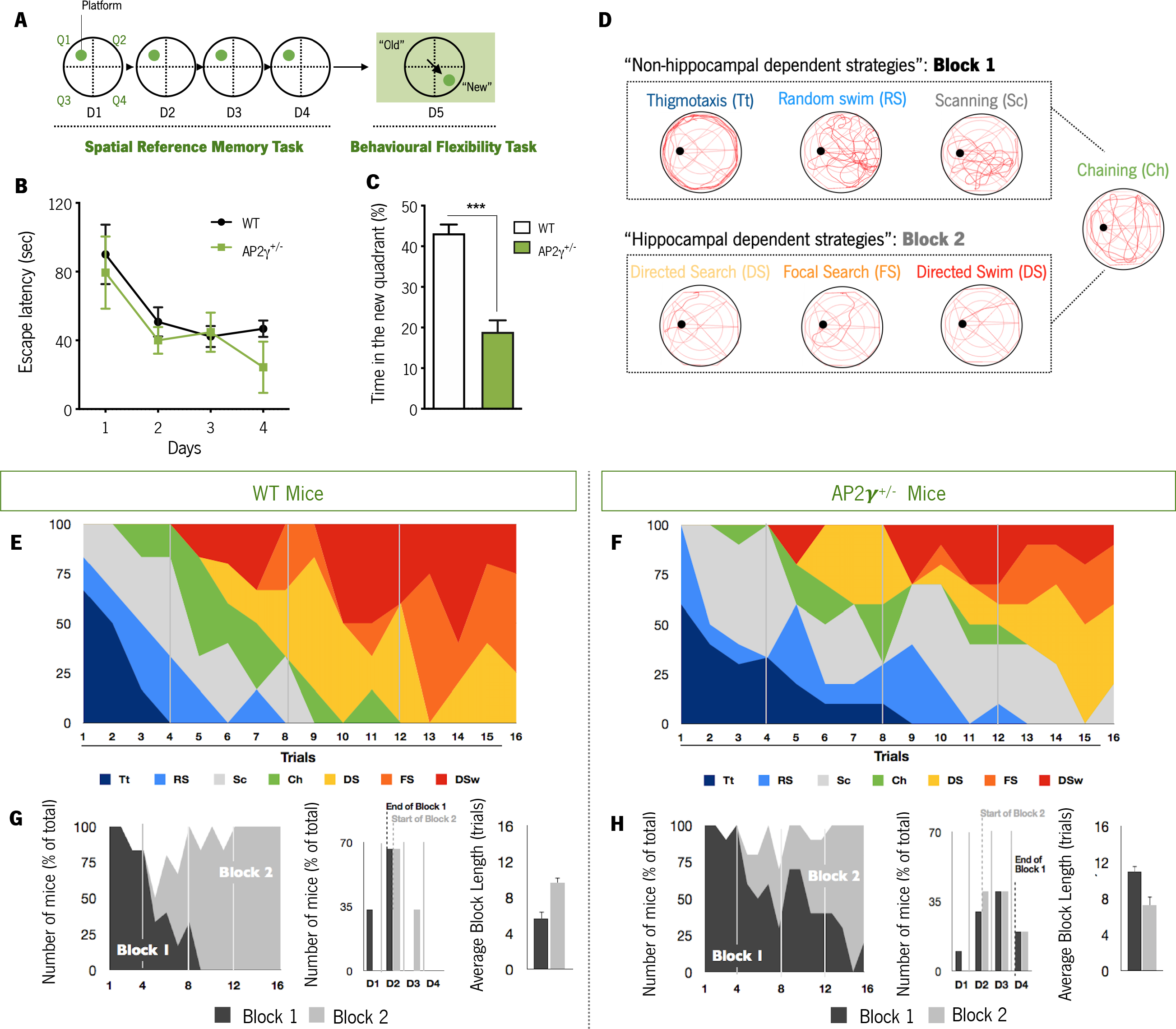
Cognitive performance of adult mice in the Morris water maze test. (A and B) Spatial reference memory was assessed as the average escape latency to find a hidden and fixed platform in each test day. (C) In the testing day, animals were subjected to a reversal-learning task to test behavioral flexibility. (D) Schematic representation of typical strategies to find the platform during spatial memory evaluation grouped according to its dependence of the hippocampus (Block 1: Non-hippocampal dependent strategies; Block 2: Hippocampal dependent strategies). Average of each strategy used for WT (E) and AP2γ^+/−^ (F) mice, by trial number. The prevalence of each block along with trials, the distribution of strategies-block boundaries, and overall block length are shown for (G) WT and (F) AP2^+/−^ mice. Data presented as mean SEM. *n*_WT_ = 10; *n*_AP2γ^+/−^_ = 10. [Repeated measures ANOVA and Student’s t-test; \*\*\**p*< 0.001; Statistical summary in Supplementary table 1]. Abbreviations: WT, wild-type; AP2γ^+/−^, AP2γ heterozygous knockout mice.

Overall, results suggest that constitutive heterozygous deletion of AP2γ resulted in specific emotional and cognitive impairments at adulthood. More specifically, we observed that adult AP2γ^+/−^ mice display anxiety-like behavior, and cognitive impairments, in contextual memory, spatial memory and behavioral flexibility. Importantly, due to the relevance of hippocampal and mPFC to these behavioral tasks, results suggest that, in adult mice, the functional integrity of these brain areas is highly affected by constitutive and heterozygous deficiency of AP2γ.

### Adult hippocampal-to-PFC functional connectivity is disrupted by constitutive AP2γ heterozygous deficiency

Given the impact of AP2γ deficiency on emotional and cognitive behavior, we sought for a functional correlate by investigating related neurocircuits. In adult WT and AP2γ^+/−^ mice, we explored the integrity of the dorsal hippocampus (dHip)-to-medial prefrontal cortex (mPFC) circuitry, assessing electrophysiological features of local field potentials (LFPs) simultaneously in these connected brain areas (Figure 5A and Supplementary Figure 5A). In AP2γ^+/−^ mice, the temporal structure of LFPs recorded simultaneously in the dHip and mPFC was affected. Specifically the spectral coherence between these regions (Adhikari et al., 2010; Oliveira et al., 2013; Sardinha et al., 2017) in AP2γ^+/−^ mice is significantly decreased in all frequency bands when compared to WT littermates (Figure 5B), indicating an impaired functional connectivity between these two brain regions in AP2γ^+/−^ mice. While in the dHip, constitutive AP2γ deficiency had a subtle impact in PSD values specifically in the Theta and Beta frequency bands (Figure 5C), in the mPFC, PSD values in all frequencies evaluated were significantly lower than WT mice (Figure 5D).

**Figure 5:**
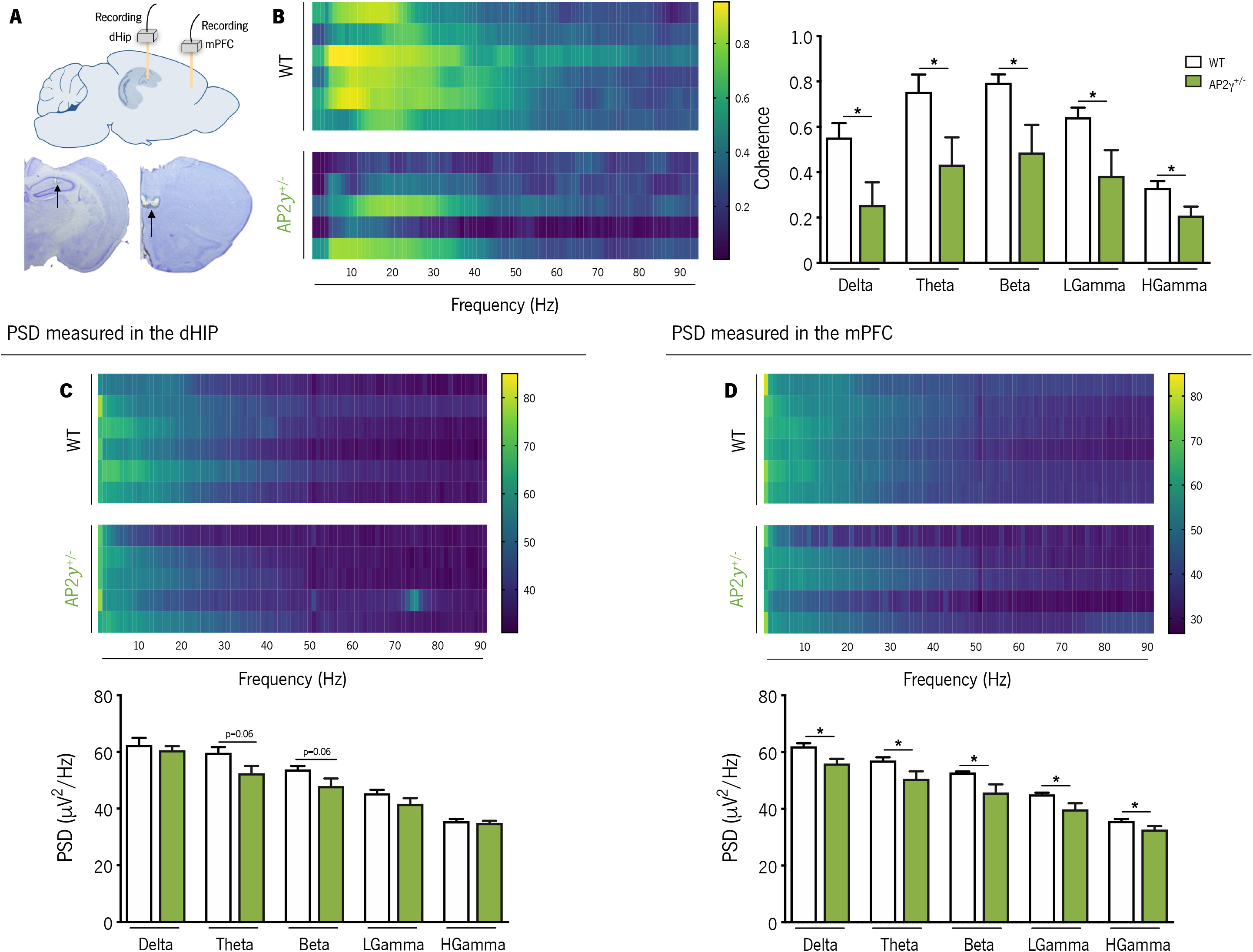
AP2γ deficiency induces deficits in spectral coherence between the dorsal hippocampus (dHip) and the medial prefrontal cortex (mPFC), impacting neuronal activity. (A) Identification of the local field potential (LFP) recording sites, with a depiction of the electrode positions (upper panel), and representative Cresyl violet-stained sections, with arrows indicating electrolytic lesions at the recording sites (lower panel). (B) Spectral coherence between the dHip and mPFC (left panel). Group comparison of the coherence values for each frequency (right panel). (C) Power spectral density (PSD) was measured in the dHip (C) and mPFC (D). Heatmaps of PSD activity (upper panel) and group comparison for each frequency (lower panel). Each horizontal line in the Y-axis of the presented spectrograms represents an individual mouse. Frequency bands range: delta (1-4Hz), theta (4-12 Hz), beta (12-20 Hz), low gamma (20-40 Hz), and High gamma (40-90 Hz). Data presented as mean SEM. *n*_WT_ = 6; *n*_AP2γ^+/−^_ = 5. [Student’s t-test; \**p*< 0.05; Statistical summary in Supplementary table 1]. Abbreviations: WT, wild-type; AP2γ^+/−^, AP2γ heterozygous knockout mice.

Deficiency of AP2γ did not exert an effect neither in the spectral coherence between the vHip and the mPFC (Supplementary Figure 5B) nor in the PSD values in the vHip (Supplementary Figure 5C).

The electrophysiological studies revealed that constitutive and heterozygous AP2γ deficiency led to two outcomes: first, a significant decrease of coherence between the dHip and the mPFC indicating impairments in the ability of these regions to functionally interact; second, this decrease in interregional coherence was accompanied by a diminished neuronal activity in a wide range of frequencies in the mPFC, including in theta and beta frequencies, previously shown to be critically related with behavioral outputs dependent on cortico-limbic networks (Colgin, 2011; Fell and Axmacher, 2011; Oliveira et al., 2013).

## Discussion

Herein, we show that despite a normal early postnatal acquisition of neurodevelopmental milestones, constitutive and heterozygous deficiency of AP2γ induces an anxiety-like state and causes cognitive deficits in mice that persist from adolescence until adulthood. Our results suggest that AP2γ plays a crucial role for the proper development and maturation of neural circuits implicated in emotional and cognitive functions.

Newly generated neurons are highly relevant to hippocampal functioning and hippocampal-associated behaviors (Anacker and Hen, 2017; Christian et al., 2014; Fang et al., 2018; Gonçalves et al., 2016). Impairments in adult hippocampal neurogenesis precipitate the emergence of depressive- and anxiety-like behaviors (Bessa et al., 2009; Hill et al., 2015; Mateus-Pinheiro et al., 2013b, 2013a; Revest et al., 2009; Sahay and Hen, 2007). Here, we assessed the longitudinal impact of AP2γ, a transcription factor that plays an important role on embryonic neuronal development (Pinto et al., 2009) and recently described as a novel regulator of adult hippocampal neurogenesis (Mateus-Pinheiro et al., 2018, 2017), on neural plasticity, function and behavior at different postnatal periods. Characterization of the neurogenic process in the hippocampal DG in juvenile and adult AP2γ^+/−^ mice, revealed that in agreement with a previous report in a conditional knock-out mice (Mateus-Pinheiro et al., 2017), AP2γ regulates upstream neurogenic regulators as Pax6 and Tbr2. Other modulators of the TAP’s population, such as Ngn2 and Tbr2, have been shown to exert a similar control of hippocampal neurogenesis. Ngn2 has been implicated in the proper development of the DG, and its deletion at early stages of development leads to a reduction in neurogenesis (Galichet et al., 2008; Roybon et al., 2009). Moreover, Tbr2 is critically required for hippocampal neurogenesis in the developing and adult mice, with its postnatal inactivation resulting in a marked reduction in neuroblasts (Hodge et al., 2012), and its conditional deletion in the adult hippocampal neurogenic niche leading to a specific blockade of the neurogenic process (Tsai et al., 2015). Also, we observed that, at both postnatal periods, AP2γ plays an essential role in the regulation of pivotal neurogenic steps as NSCs proliferation and neuronal maturation (Mateus-Pinheiro et al., 2017), while the morphology of granular neurons in the hippocampal DG, another form of hippocampal structural plasticity, is intact (Bessa et al., 2009; Mateus-Pinheiro et al., 2013a).

Taking into consideration the embryonic and early postnatal developmental modulatory roles of AP2γ, and the severe and/or lethal malformations during development promoted by deficiencies in other members of the AP2 family (AP2α and AP2β) (Lim et al., 2005; Moser et al., 1997; Schorle et al., 1996), we sought to understand whether constitutive and heterozygous deficiency of AP2γ could lead postnatally to functional and behavioral impairments. The developmental milestones protocol showed no impact of the constitutive heterozygous deficiency of AP2γ in early postnatal neurodevelopment. Nevertheless, this deficiency in AP2γ promotes emotional and cognitive behavior in later periods of life. At juvenile age, AP2γ deficiency led to the manifestation of anxiety-like behavior and significant impairments in recognition memory tasks, that depend on the integrity of the hippocampal circuitry (Jessberger et al., 2009). Interestingly, anxiety-like behavior and cognitive impairments were maintained at adulthood, where adult AP2γ^+/−^ mice displayed poor performances in hippocampal-dependent tasks (Garthe et al., 2009; Garthe and Kempermann, 2013; Gu et al., 2012; Ruediger et al., 2012). Notably, conditional deletion of AP2γ in adulthood lead to a less evident effect on emotional behavior, namely in anxiety-like behavior tested in the OF and EPM behavioral tests, when comparing with the constitutive mice model herein presented (Mateus-Pinheiro et al., 2017). This result indicates that constitutive deficiency of AP2γ may exert a longitudinal cumulative impact leading to more severe alterations in behavioral performance of mice, whereas in the conditional model during adulthood, the AP2γ deletion only occurs in a subset of newly formed neuroblasts.

Our results are consistent with previous publications were the suppression of the TAP’s regulator Tbr2 exerted both an anxiety-like phenotype during the juvenile period, and induced cognitive deficits during early adulthood (Veerasammy et al., 2020). Moreover, Ngn2 is also important for the modulation of cognitive behavior, namely in the rescue of cognitive function in the T-Maze task. Interestingly, the regulation of TAPs’ by AP2γ seems to be also important for the preservation of cognitive performance as shown by its impact on hippocampal-dependent tasks. Behavioral flexibility, a cognitive task that relies on the interaction of the hippocampal and prefrontal cortical brain areas, was impaired in AP2γ^+/−^ mice. Adult AP2γ^+/−^ mice present significant deficits of electrophysiological coherence between the dHip and the mPFC. In particular, constitutive AP2γ deficiency led to a decrease of the spectral coherence between the recorded brain areas in a wide range of frequencies, previously associated to behavior outputs dependent on cortico-limbic networks (Colgin, 2011; Fell and Axmacher, 2011; Oliveira et al., 2013; Sardinha et al., 2017). The integrity of the hippocampus-to-PFC circuitry was described to be relevant for example to the action of antidepressants, such as ketamine (Carreno et al., 2016), which promote neurogenesis, suggesting that AP2γ may be involved in conserving this neuronal circuit. Additionally, AP2γ plays an important role on cortical basal progenitors’ specification during embryonic development (Pinto et al., 2009) that might be affecting the electrophysiological function of the mPFC. In fact, AP2γ^+/−^ mice presented impaired neuronal activity in the mPFC in all frequency ranges, as detected by the general decrease of PSD signals recorded, and corroborated by previous findings (Mateus-Pinheiro et al., 2017). Thus, misspecification of upper cortical layers promoted by AP2γ deficiency since embryonic development may be contributing to the functional electrophysiological readouts, and also eliciting the cognitive defects herein observed.

Collectively, the results presented in this study demonstrated the importance of the transcription factor AP2γ in the generation of glutamatergic neurons in the postnatal brain and its impact on functional behavioral dimensions at different postnatal periods. Following these findings, future experiments should be implemented to elucidate whether AP2γ can participate in the pathogenesis and treatment of neurodevelopmental and/or psychiatric disorders and how it may exert its modulatory action.

## Materials and methods

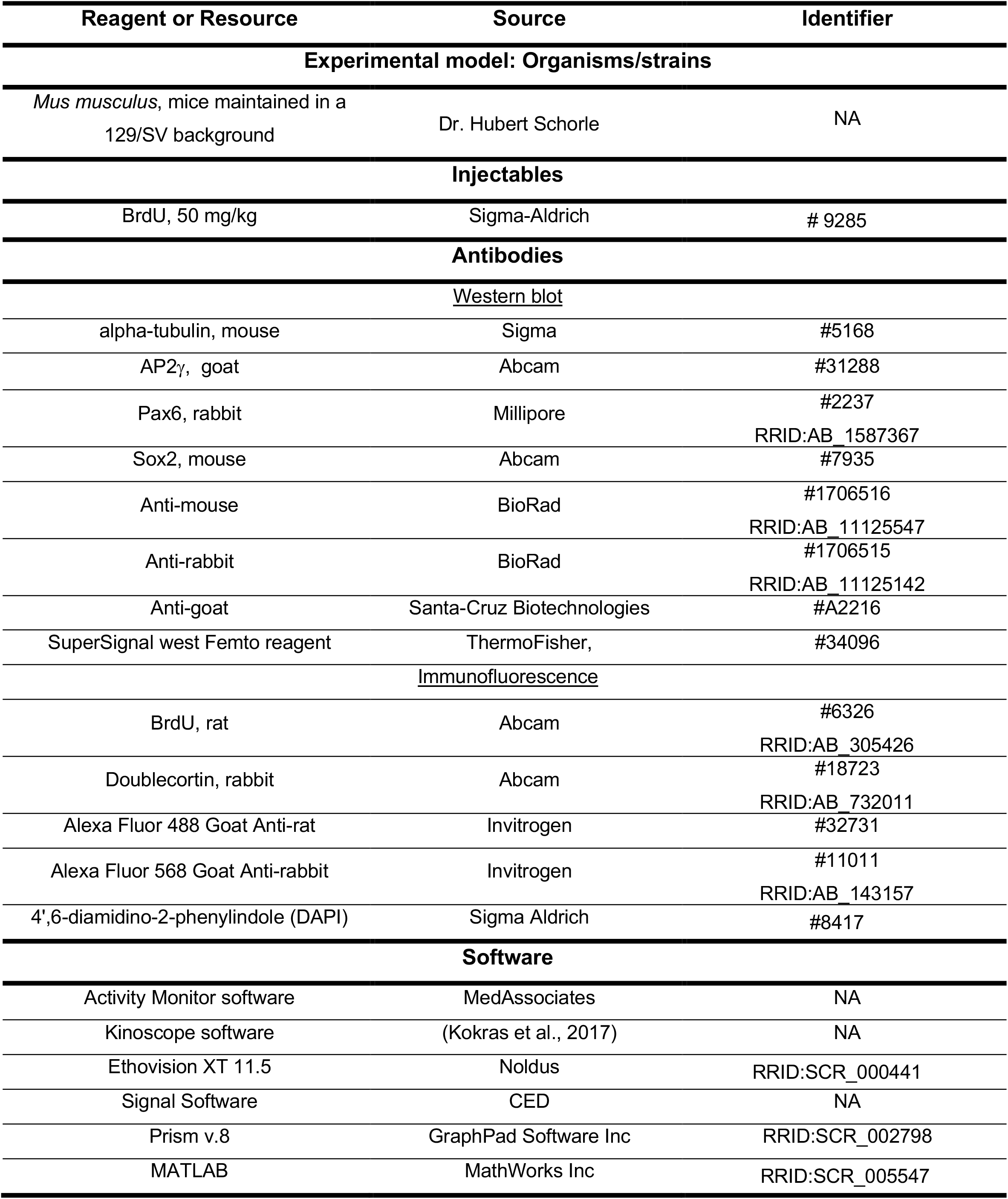
Key Resource Table.

### Experimental model details

Wild-type (WT) and AP2γ heterozygous KO (AP2γ^+/−^) mice were maintained in a 129/SV background and identified by polymerase chain reaction (PCR) of genomic DNA. Along the study, distinctive at cohorts (at least 2 *per* timepoint) of littermate WT and AP2γ^+/−^ male mice were submitted to molecular (n = 4-6 *per* group), behavioral (n = 8-16 *per* group) and electrophysiological (n= 5-6 *per* group) assessment at the different postnatal ages (early postnatal period: between postnatal day (PND) 1 and 21; juvenile period: from PND 25 to 31; adulthood: PND 70 to 92).

All mice were housed and kept under standard laboratory conditions at 22 ± 1°C, 55% humidity, and *ad libitum* access to food and water on a 12h light/dark cycle (lights on 8 A.M. to 8 P.M.). Efforts were made to minimize the number of animals and their suffering. All experimental procedures performed in this work were conducted in accordance with the EU Directive 2010/63/EU and approved by the Portuguese National Authority for animal experimentation, *Direção-Geral de Alimentação e Veterinária* (DGAV) with the project reference 0420/000/000/2011 (DGAV 4542).

### Behavioural analysis

#### Developmental milestones protocol

Early postnatal neurodevelopment in mice was assessed according to previously validated protocols (Castelhano-Carlos et al., 2010; Guerra-Gomes et al., 2020; Hill et al., 2008; Santos et al., 2007). This consisted in a daily evaluation for the first 21 days of life. From postnatal day (PND) 1 onward, newborn animals were evaluated in several parameters, including skin appearance, activity, and presence of milk spot in the stomach, indicator of correct maternal care and well-being. Pups were examined for the acquisition of developmental milestones until weaning (PND21), every day at the same time, in the same experimental room, by the same experimenter. This daily scoring included tests to assess the acquisition of mature response regarding somatic parameters and neurobiological reflexes.

##### Somatic parameters

As a measure of morphological development, animals were daily weighed (weight ± 0.01 g). The eye-opening day was also evaluated and considered when both eyes were opened. When both eyes opened on different days, score was set as 1 if only one of the eyes was open, and 2 when both eyes were open. The mature response was registered when both eyes were open.

##### Neurobiological reflexes

The assessment of the neurobiological reflexes including the daily performance of different tests. Of note, the scale of evaluation was distinct among tests. Tests including rooting, ear twitch, auditory startle, open field transversal, air righting, wire suspension, postural reflex were scored according to the absence (0) or presence (1) of a mature response. When possible, to detect a gradual progression in performance as for walking, surface righting, grasping, negative geotaxis, cliff aversion, daily score was attributed between 0 and 3, with 0 representing absence, and 3 corresponding to the achievement of the mature response. The postnatal day in which animals achieved a mature response was registered. All tests were conducted in a smooth foam pad, and immediately after testing, the pups were returned to their home cage.

##### Labyrinthine reflex, body righting mechanism, coordination and strength

*Surface righting reflex* – PND1 to PND13 – This test consists of gently laying the animal on its back, and the mature response was considered when the pup was able to get right. If the animal did not respond within 30 s, the test was ended. Mature response was achieved when the pups were able to get right in less than 1 s for three consecutive days.

*Negative geotaxis* – PND1 to PND 14 – Pups were placed head down in a horizontal grid, tilted 45° to the plane. The acquisition of a mature response was set when pups were able to head up in less than 30 s for three consecutive days.

*Air righting* – PND8 to PND 21 – In this test, the pup was held upside down and released from a height of approximately 13 cm from the soft padded surface and released. A mature response was obtained when the animals landed on four paws for three consecutive days.

*Cliff aversion* – PND 1 to PND 14 – It evaluates the mouse pup’s ability to turn and crawl away when on the edge of a cliff. A mature response was achieved once the animal moved away in less than 30 s for three consecutive days.

*Postural reflex* – PND 5 to PND 21 – Pups were placed in a small plastic box and gently shaken up down and right. When the animals were able to maintain their original position in the box by extending four paws, mature response was acquired.

*Wire suspension* – PND 5 to PND 21 – This test evaluates forelimb grasp and strength. Pups were placed vertically to hold with their forepaws a 3 mm diameter metal wire suspended 5 cm above a soft foam pad. A mature response was achieved once the animal was able to grasp the bar, holding it with four paws.

*Grasping* – PND 5 to PND 21 – The mouse pup forelimb was stimulated with a thin wire to evaluate when the involuntary freeing reflex stopped. This reflex disappears with the development of the nervous system, as so, the mature response achieved when the animal grasped immediately and firmly the wire.

##### Tactile reflex

*Ear twitch* – PND 7 to PND 15 – In this test, the mouse pup ear was gently stimulated with the tip of a cotton swab, three times. If the animal reacted, flattening the ear against the side of the head for three consecutive days, the mature response was reached.

*Rooting* – PND 7 to PND 12 – A fine filament of a cotton swab was used to gently and slowly rub the animal’s head, from the front to the back. It was considered a successful test if the pup moved its head towards the filament. The test was repeated on the other side of the head to evaluate the appearance of this neurobiological reflexes on both sides. If the animal did not react to the filament, the test was repeated. Mature response was obtained when animal reacted on both sides for 3 consecutive days.

##### Auditory reflex

*Auditory startle* – PND7 to PND 18 – We evaluated the reaction of pups to a handclap, at a distance of 10 cm. If pups quick and involuntary jumped for three consecutive days, a matured response was attributed.

##### Motor

*Open field transversal* – PND7 to PND 18 – To execute this test, animals were placed in a small and circle (13 cm diameter), and time to move was recorded. If the pup was not able to move, the test was ended. In case the mouse leaves the circle in less than 30 s, in three consecutive days, a mature response was reached.

*Walking* – PND 5 to PND 21 – In this paradigm, animals were able to freely move around for 60 s. The mature response was achieved when they showed a walking movement fully supported on 4 limbs.

### Open field (OF) test

The OF test is a behavioral test commonly used to assess anxiety-like behavior, exploratory behavior, and general activity in rodents. This apparatus consists of a highly illuminated square arena of 43.2×43.2cm, closed by a 30.5 cm high wall. Mice were individually placed in the center of the OF arena, and their movement was tracked for 5 mins, using a 16-beam infrared system (MedAssociates, US). Data was analysed using the Activity Monitor software (MedAssociates, US). Average velocity of the animals was considered as a measure of locomotor capacity, and the activity ratio between the center and the arena periphery was considered as measurement of anxiety-like behavior.

### Elevated-plus maze (EPM) test

To study the impact on anxiety-like behavior the EPM test was also performed (Walf and Frye, 2007). This consists of a black propylene apparatus (ENV – 560; MedAssociates Inc, US) with two opposite open arms (50.8 cm × 10.2 cm) and two closed arms (50.8 cm × 10.2 cm × 40.6 cm) elevated 72.4 cm above the floor and dimly illuminated. The central area connecting both arms measured 10×10 cm. Animals were individually positioned in the center of the maze, facing an edge of a closed-arm, and were allowed to freely explore the maze for 5 min. All trials were recorded using an infrared photobeam system, and the percentage of time spent in the open arms was accessed through the EthoVision XT 11.5 tracking system (Ethovision, Noldus Information Technologies, Netherlands).

### Forced swimming test (FST)

For the assessment of behavioral-despair, we performed the forced swimming test (Porsolt et al., 1977). Briefly, each animal was individually placed in glass cylinders filled with water (23°C; depth 30 cm) for 5 min. All sessions were video-recorded, and the immobility time, defined through a video tracking software Ethovision XT 11.5 (Noldus, Netherlands), was considered as a measure of learned-helplessness. Mice were considered immobile when all active behaviors (struggling, swimming, and jumping) were ceased. For immobility, the animals had to remain passively floating or making minimal movements need to maintain the nostrils above water. For learned-helplessness assessment, the first 3 min of the trial were considered as a habituation period and the last 2 min as the test period.

### Tail-suspension test (TST)

The TST is a commonly used behavioral test to assess behavioral despair in rodents. The principle of this behavioral paradigm is similar to the FST assessing also learned helplessness of the animals. For this, animals were suspended by the tail to the edge of a laboratory bench 80 cm above the floor (using adhesive tape) for 6 min. Trials were video-recorded, and the immobility and climbing times were automatically analyzed by the video tracking software Ethovision XT 11.5 (Noldus, Netherlands). For the learned-helplessness assessment, the first 3 min of the trial was considered as a habituation period, and the last 3 min as the test period.

### Splash-sucrose test (SCT)

The splash-sucrose test consists of spraying a 10 % sucrose solution on the dorsal coat of mice in their home cage (Yalcin et al., 2008). Sucrose solution in the mice’s coat induces grooming behavior. After spraying the animals, animal was video-recorded for 5 min and the time spent grooming was taken as an index of self-care and motivational behavior. Then, the videos were manually analyzed using the behavioral scoring program Kinoscope (Kokras et al., 2017).

### Object recognition test (ORT)

Through this behavioral paradigm we assessed short- and long-term memory (Leger et al., 2013). This test relies on rodents’ nature to explore and prefer novelty. For that, mice were acclimatized to a testing arena (30 cm × 30 cm × 30 cm) under dim light for 3 days during 20 min. After habituation, animals were presented with two equal objects for 10 min (training), positioned in the center of the arena. Then, 1 h later, one of the objects was moved towards one arena wall, and mice were allowed to freely explore the objects for 10 mins. On the following day, animals returned to the arena for 10 mins, with one of the objects replaced by a novel object. The familiar and novel objects had different size, color, shape and texture. Between trials, the arena and objects were properly cleaned with 10% ethanol. Sessions were recorded and manually scored through the behavioral scoring program Kinoscope (Kokras et al., 2017). The percentage of time exploring the moved- and novel-object was used as a measure of short- and long-term memory, respectively.

### Contextual-fear conditioning (CFC)

The CFC test was performed in a white acrylic box with internal dimensions of 20 cm wide, 16 cm deep, and 20.5 cm high (MedAssociates). This apparatus had a fixed light bulb mounted directly above the chamber to provide a source of illumination. Each box contained a stainless-steel shock grid floor inside a clear acrylic cylinder, where the animals were placed. All animals were exposed to two probes: a context probe and a cue (light) probe, as previously described (Gu et al., 2012; Mateus-Pinheiro et al., 2017). All probes were recorded, and the freezing behavior was manually scored through Kinoscope (Kokras et al., 2017). This behavioral paradigm took 3 days.

#### Day 1

Animals were individually placed in the conditioning-white box (Context A) and received three pairings between a light (20 s) and a co-terminating shock (1 s, » 0.5 mA). The interval between pairings was 180 s, and the first light presentation started 180 s after the beginning of the trial. After the three pairings, mice remained in the acrylic box for 30 s, being after returned to their home cage. Between animals, the apparatus was properly cleaned with 10% ethanol.

#### Day 2

For the context probe, animals were placed into the same white acrylic chamber (context A), 24h hours after the light-shock pairings. The freezing behavior was monitored for 3 min. Two hours later, we introduced the animals into a modified version of the chamber (Context B). This new box was sheeted with a black plasticized cover, sprayed with a vanilla scent. In this way, both contexts had distinct spatial and odor cues. Also in Context B, the ventilation was not operated, and the experimenter wore a different color of gloves and a lab coat. Freezing behavior was measured for 3 min. The freezing behavior state was defined as the total absence of motion, for a minimum of 1 s.

#### Day 3

For the cue probe, the animals were set in Context B, and individually placed in this chamber 24h after the context probe. After 3 min, the light was turned on for 20 s, and the freezing behavior monitored for 1 min after light is turned off.

### Morris water maze (MWM)

In the MWM test, several cognitive domains were assessed: working- and spatial-reference memory and behavioral flexibility. Additionally, the strategies used to reach the platform were also analyzed. MWM was performed in a circular white pool (170 cm diameter) filled with water at 22°C to a depth of 31 cm in a room with and dim light and extrinsic clues (triangle, square, cross, and horizontal stripes). The pool was divided into four quadrants by imaginary lines, and a clear-acrylic cylinder platform (12 cm diameter; 30 cm high), placed in one of the quadrants. All trials were video recorded by a tracking system (Viewpoint, France).

#### Working memory task

The working memory task (Alves et al., 2017; Cerqueira et al., 2007) evaluates the cognitive domain that relies on the interplay between the hippocampal and prefrontal cortex (PFC) functions. In this task, animals had to learn the position of the hidden platform and to retain this information for four consecutive daily trials. The task was performed during 4 days and in a clockwise manner the platform was repositioned in a new quadrant each day. During the daily trials, animals had different starting positions (north, east, west, and south). Trials ended when the platform was reached within the time limit of 120 s. If the animals did not reach the platform during the trial time, they were guided to the platform and allowed to stay for 30 s. The time and path to reach the platform were recorded.

#### Reference memory task

After working memory evaluation (days 1-4), spatial-reference memory, a hippocampal dependent-function, was assessed by keeping the platform in the same quadrant during three consecutive days (days 4-6) (Morris, 1984). The time and path to reach the platform were recorded for each trial.

#### Reversal learning task

On the last day of MWM testing, reversal-learning performance, a PFC dependent function, was assessed. This was conducted by positioning the platform in a new (opposite) quadrant. Animals were tested in 4 trials. The percentage of time spent in the new and old quadrant containing the platform was used as readout of behavioral flexibility.

#### Search strategies analysis

Throughout the Morris water maze, animals were evaluated through the adopted strategies to reach the hidden platform, as previously described (Garthe and Kempermann, 2013; Mateus-Pinheiro et al., 2017; Ruediger et al., 2012). Quantitative analyses and strategy classification were completed by assessing different parameters collected through the Viewpoint software: (1) thigmotaxis (Tt): most of the swim distance (>70%) happened within the outer ring area (8 cm from the pool border; (2) random swim (RS): most of the swim distance (>80%) occurred within the inner circular area, and all quadrants were explored with a percentage of swim distance not below 50% for none of the quadrants; non-circular trajectories; (3) scanning (Sc): most of the swim pattern and distance (>80%) happened within the inner circular area, with balanced exploration in all quadrants of the pool; non-circular trajectories, with a percentage (<60%) of swim distance in the platform corridor area (area centered along the axis that connects the start position and the hidden platform); (4) chaining (Ch): the majority of the swim distance occurred in the inner circular area (>80%), with a balanced exploration of all pool quadrants; swim distance in the platform corridor area <60%, with circular trajectories taking place; (5) directed search (DS): the majority of the swim distance occurred in the inner circular area (>80%); swim distance in the platform corridor area >60%, with shifts in the trajectories directions; (6) focal search (FS): directed trajectories to the platform zone, with swim exploration within the perimeter of the escape platform (30cm); (7) directed swim (DSw): directed trajectories to the hidden platform, without much exploration of the pool. For simplification, we defined two blocks of strategies: Block 1, that comprises the “non-hippocampal dependent strategies” (Tt, RS, and Sc), and Block 2, comprising the defined “hippocampal dependent strategies” (DS, FS, and DSw). These blocks were defined when a sequence of at least three trials within the same block were reached.

### Electrophysiological studies

Electrophysiological recordings were obtained from anesthetized mice (sevoflurane 2,5%; 800 mL/min). A surgical procedure was performed to insert platinum/iridium concentric electrodes (Science Products) in the target positions following the mouse brain atlas (from Paxinos): prelimbic region of the medial prefrontal cortex (mPFC): 1.94 mm anterior to bregma, 0.4 mm lateral to the midline, 2.5 mm below bregma; dorsal hippocampus (dHIP): 1.94 mm posterior to bregma, 1.2 mm lateral to the midline, 1.35 mm below bregma); ventral hippocampus (vHIP): 3.8 mm posterior to bregma, 3.3 mm lateral to the midline, 3.4 mm below bregma). LFP signals obtained from mPFC, dHIP, and vHIP were amplified, filtered (0.1–300 Hz, LP511 Grass Amplifier, Astro-Med), acquired (Micro 1401 mkII, CED) and recorded through the Signal Software (CED). Local field activity was recorded at the sampling rate of 1000 Hz during 100s. After electrophysiological recordings, a biphasic 0.7 mA stimulus was delivered to mark the recording sites. Then, mice were deeply anesthetized with sodium pentobarbital, brains removed, immersed in paraformaldehyde (PFA) 4% for 48h and sectioned (50 μm) in the vibratome. Coronal slices containing the mPFC, dHip vHip were stained for Cresyl Violet to check for recording sites. Animals with recording positions outside at least in one of the two regions under study (mPFC and dHip or vHip) were excluded from the analysis. Coherence analysis was based on multi-taper Fourier analysis.

Coherence was calculated by custom-written MATLAB scripts, using the MATLAB toolbox Chronux (http://www.chronux.org) (Mitra and Pesaran, 1999). Coherence was calculated for each 1 s long segments and their mean was evaluated for all frequencies from 1 to 90 Hz. The power spectral density (PSD) of each channel was calculated through the 10 × log of the multiplication between the complex Fourier Transform of each 1s long data segment and its complex conjugate. The mean PSD of each channel was evaluated for all frequencies from 1 to 90 Hz (Oliveira et al., 2013). Both coherence and PSD measurements were assessed in the following frequencies: delta (1–4 Hz), theta (4–12 Hz), beta (12–20 Hz); low (20–40 Hz) and high gamma (40-90 Hz).

### BrdU labelling

To assess the effect of AP2γ heterozygous deletion on the proliferation of fast-dividing progenitor cells, and its impact on the generation of adult-born neurons, animals from all groups were injected intraperitoneally with the thymidine analogous 5-bromo-2’-deoxyuridine or bromodeoxyuridine (BrdU, 50 mg/kg; Sigma-Aldrich, US) that is incorporated in the DNA during the S-phase. BrdU injections were performed once, at the end of the behavioral assessment, 24 h prior to occision.

### Western Blot analysis

Hippocampal DG of juvenile and adult AP2γ^+/−^ mice and WT littermates were carefully macrodissected out after occision. The tissue was weighted and homogenized in RIPA buffer [containing 50mM Tris HCl, 2 mM EDTA, 250 mM NaCl, 10 % glycerol, 1 mM PMSF protease inhibitors (Roche, Switzerland)] and then sonicated (Sonics & Materials, US) for 2 min. Samples were centrifuged for 25 min at 10.000 rpm and 4°C. The protein concentration of the supernatant was determined using Bradford assay. Samples with equal amounts of protein, 30 μg, were analyzed using the following primary antibodies: alpha-tubulin (#5168; Sigma, mouse, 1:5000), AP2γ (#31288; goat, 1:500; Abcam, UK), Pax6 (#2237; rabbit, 1:1000; Millipore, US), Sox2 (#7935; mouse, 1:500; Abcam, UK) and Tbr2 (#2283; rabbit, 1:500; Millipore, US). Secondary antibodies were used from BioRad (Anti-mouse, 1:10.000; #1706516; Anti-rabbit, 1:10.000; #1706515, US) and Santa-Cruz Biotechnologies (Anti-goat, 1:7500; #A2216, US). Membranes were developed using SuperSignal west Femto reagent (#34096; ThermoFisher, US) and developed in Sapphire Biomolecular Imager from Azure Biosystems (US). After developing, images were quantified using AzureSpot analysis software (Azure Biosystems, US).

### Immunostaining procedures

All mice were deeply anesthetized and then transcardially perfused with cold 0.9% NaCl, followed by 4% paraformaldehyde (PFA). Brains were carefully removed from the skull, postfixed in 4% PFA, and then cryoprotected in 30% sucrose solution. The brains were coronally processed at the vibratome (Leica VT 1000S, Germany) with a thickness of 50 mm, extending over the entire length of the hippocampal formation. Coronal sections containing the hippocampal dentate gyrus (DG) were further stained to assess cell proliferation and the population of neuroblasts. For that purpose, brain sections were double stained for BrdU (#6326; rat, 1:100; Abcam, UK) and doublecortin (DCX; #18723; rabbit, 1:100; Abcam, UK). Appropriate secondary fluorescent antibodies were used (Alexa Fluor 488 Goat Anti-rat, #32731; 1:1000; Invitrogen, US; and Alexa Fluor 568 Goat Anti-rabbit, #11011; 1:1000; Invitrogen, US). For Cell nuclei labeling, 4’,6-diamidino-2-phenylindole (DAPI, 1:200; Sigma Aldrich) was used. The density of each cell population in the DG was determined by normalizing positive cells with the corresponding area. Analysis and cell counting were performed using a confocal microscope (Olympus FluoViewTM FV1000, Hamburg, Germany) and an optical microscope (Olympus BX51). The observer was blind to the experimental condition of each subject. Data are reported as the number of cells *per* 100 μm^2^.

### 3D morphological analysis

To evaluate the 3D dendritic morphology of pre-existing granule neurons in the DG we performed impregnation with Golgi-Cox technique in brain sections from juvenile and adult mice. Briefly, brains were immersed in Golgi-Cox solution for 14 days and then transferred to 30% sucrose. Coronal sections (200 μm) were cut on a vibratome (Leica VT100S, Germany), collected and then blotted dry onto gelatine-coated microscope slides. Sections containing the dorsal hippocampus were then alkalinized in 18.8% ammonia, developed in Dektol (Kodak, US), fixed in Kodak Rapid Fix, dehydrated and xylene cleared. Dendritic arborization was analyzed in the DG of WT and AP2γ^+/−^ animals (10 neurons *per* animal).

### Data analysis and statistics

Statistical analysis was performed using Prism v.8 (GraphPad Software, US). Animals were randomly assigned to groups, balanced by genotypes. Sample sizes were determined by power analyses based on previously published studies (Mateus-Pinheiro et al., 2017) and normal distributions were assessed using the Shapiro-Wilk statistical test, taking into account the respective histograms and measures of skewness and kurtosis. To variables that followed the Gaussian distribution within groups, parametric tests were applied, while non-parametric tests were used for discrete variables. To compare the mean values for two groups, a two-tailed independent-sample t-test was applied. For comparisons between two time-points a two-way ANOVA was used. For longitudinal analyses (across days and different trials) a repeated measures ANOVA was used.

For the comparison of categorical variables (strength to grab, limb grasping and clasping), crosstabulations were performed and the statistical test used was Fisher’s exact test (when Pearson Qui-Squared assumptions were not met).

Data is expressed either as mean ± SEM (standard error of the mean), as median, or as percentage, as stated in the figures’ legends. Statistical significance was set when *p* < 0.05.

## Acknowledgments and funding

E.L.C., N.D.A., A.M.P., P.P., C.S.C., J.S., T.S.R., B.M.P., J.F.O., and L.P. received fellowships from the Portuguese Foundation for Science and Technology (FCT) (IF/00328/2015 to J.F.O.; 2020.02855.CEECIND to LP). This work was funded by FCT (IF/01079/2014, PTDC/MED-NEU/31417/2017 Grant to JFO), BIAL Foundation Grants (037/18 to J.F.O. and 427/14 to L.P.) and Nature Research Award for Driving Global Impact - 2019 Brain Sciences (to L.P.). This was also co-funded by the Life and Health Sciences Research Institute (ICVS), and by FEDER, through the Competitiveness Internationalization Operational Program (POCI), and by National funds, through the Foundation for Science and Technology (FCT) - project UIDB/50026/2020 and UIDP/50026/2020. Moreover, this work has been funded by ICVS Scientific Microscopy Platform, member of the national infrastructure PPBI - Portuguese Platform of Bioimaging (PPBI-POCI-01-0145-FEDER-022122; by National funds, through the Foundation for Science and Technology (FCT) - project UIDB/50026/2020 and UIDP/50026/2020.

## Author contributions

E.L.C. and N.D.A. maintained the AP2γ^+/−^ colony, genotyping and conducted all behavioral tests, molecular and immunohistological analyses. E.L.C. and N.D.A. also completed all the analyses and interpreted the results. P.P. assisted in the occision of the animals and performed the macrodissection of all analyzed brain areas. C.S.C. and J.S assisted in the western blots. A.M.P. helped with cognitive assessment and analyses. E.L.C., N.D.A., C.S.C., and V.M.S. collected and analyzed the electrophysiology results. T.S.R. assisted colony maintenance and genotyping. B.M.P conducted and helped with cognitive assessment and analysis. J.F.O. interpreted the electrophysiological data. E.L.C., N.D.A., and L.P. designed the study, planned the experiments, and wrote the manuscript. E.L.C., N.D.A., A.J.R., J.F.O., N.S. and L.P. edited the manuscript.

---

**Table 1:**
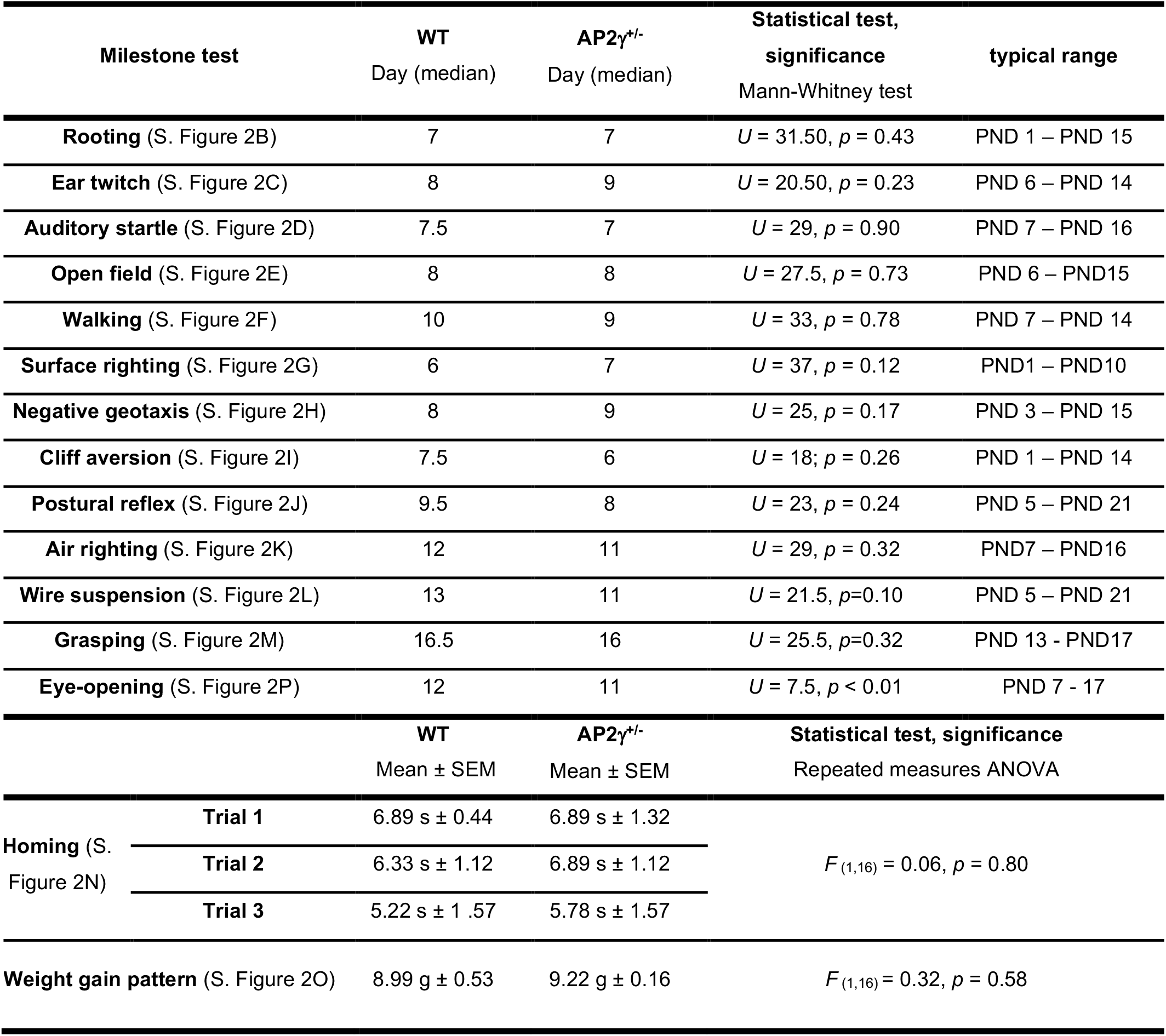
Results from the milestones protocol tests included in the assessment of early postnatal neurodevelopment. Sample size: *n*_WT_ = 9; *n*_AP2γ^+/−^_ = 9. Abbreviations: WT, wild-type; AP2γ^+/−^, AP2γ heterozygous mice; PND: Postnatal day

**Supplementary Table 1:**
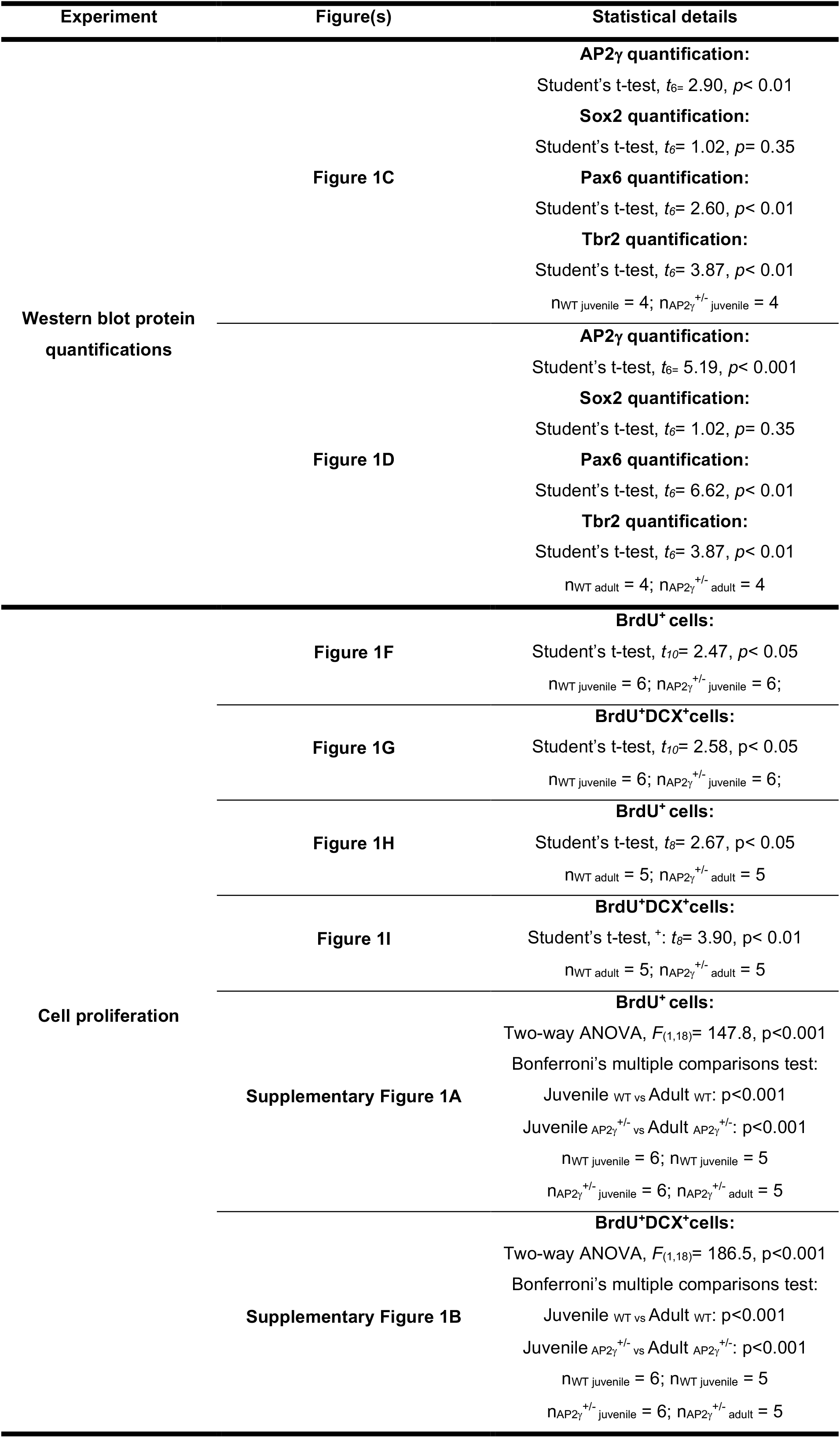
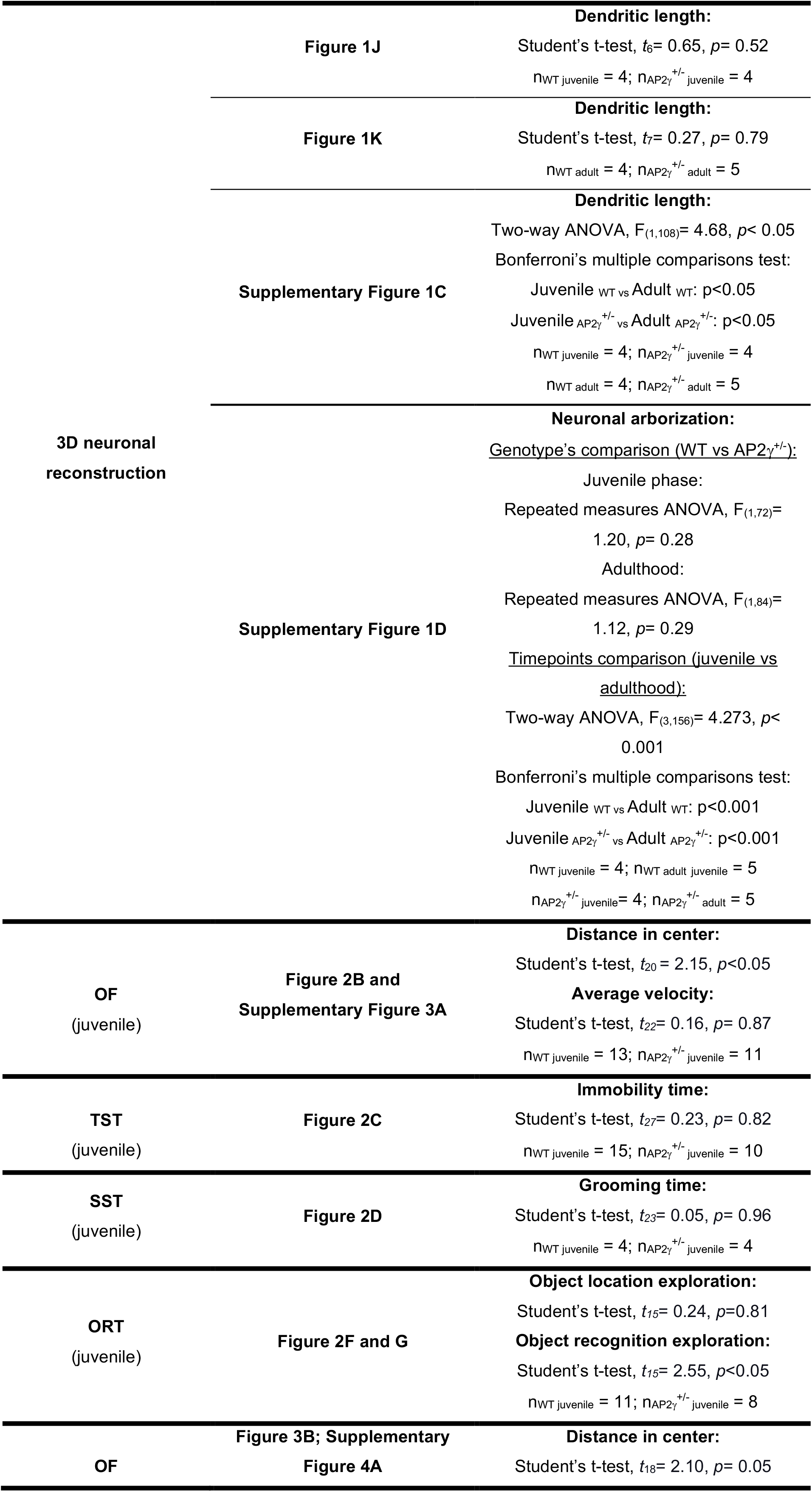
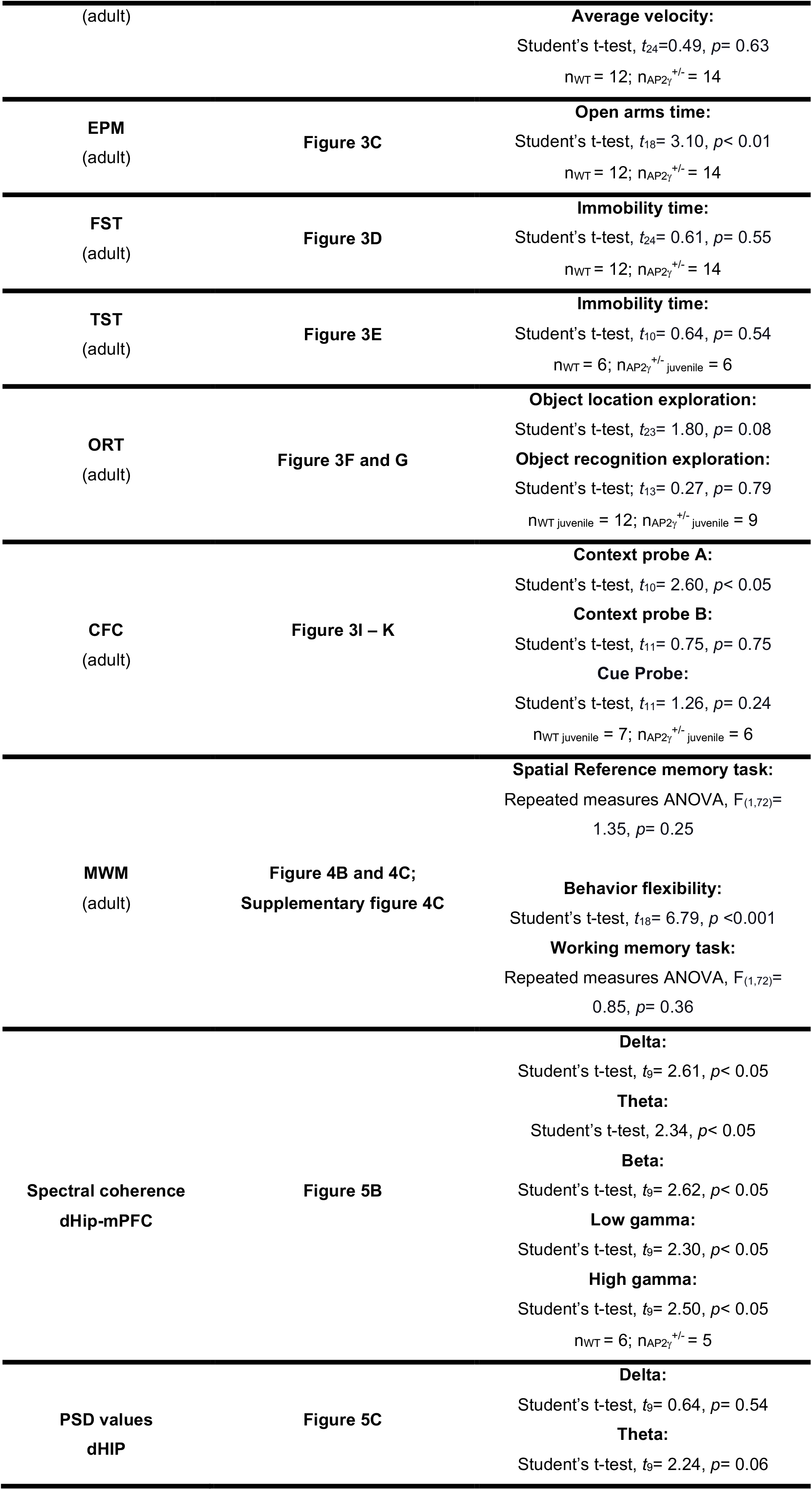
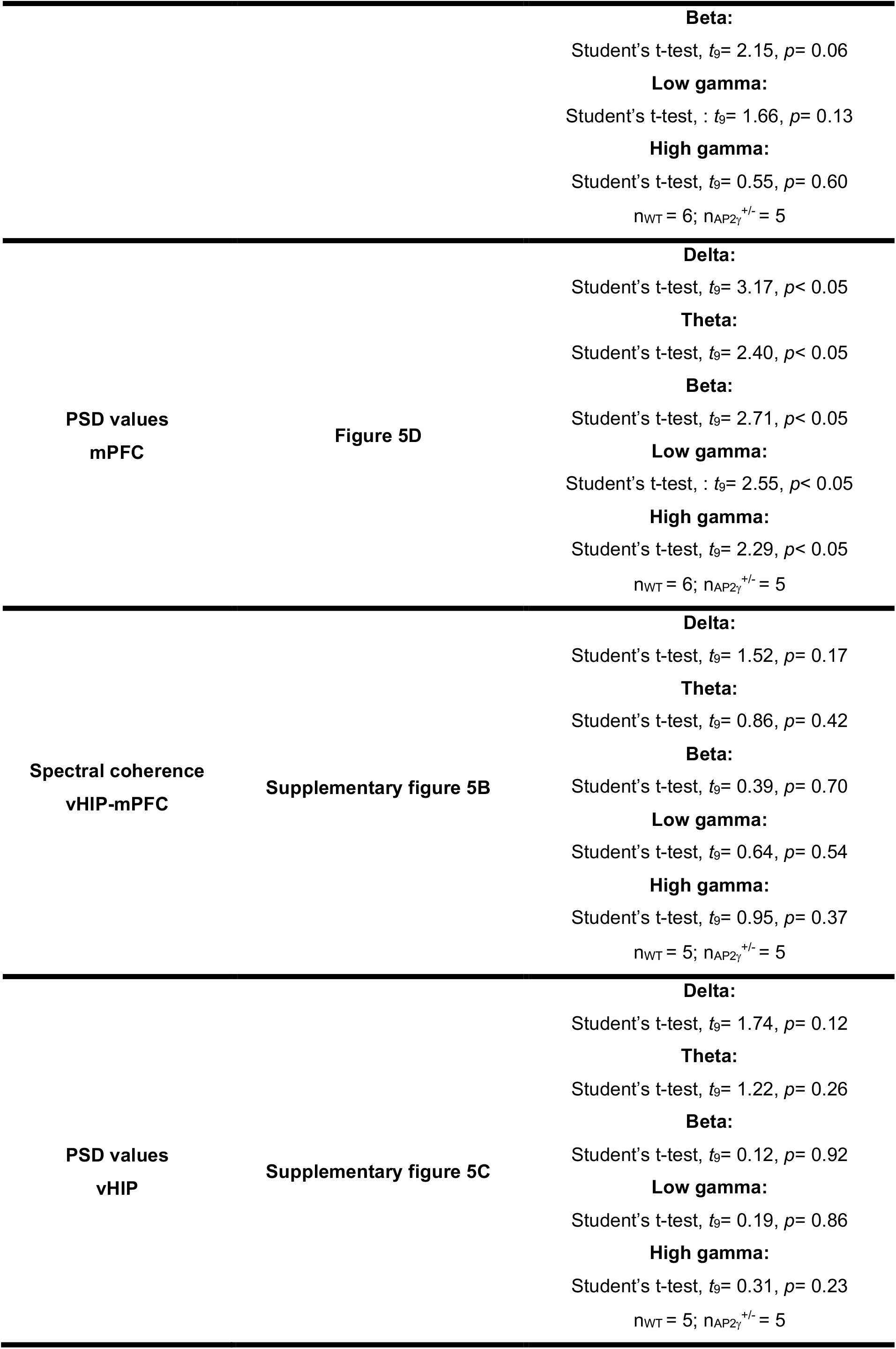
Statistical summary of results

**Supplementary figure 1:**
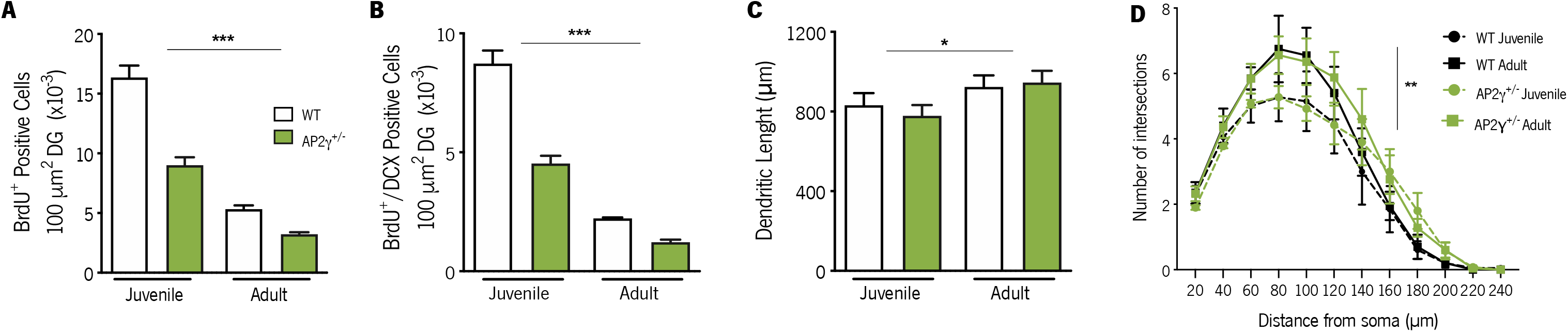
Comparative analysis of neural stem cells (NSC) proliferation and maturation in the hippocampal dentate gyrus (DG) at juvenile and adult periods. Cell counts of BrdU positive (A) and BrdU/DCX double-positive (B) cells. Dendritic length (C) and complexity (D) of hippocampal granular neurons in juvenile and adult mice. Data presented as mean SEM. Sample size: Immunostainings assays: n_WT juvenile_ = 6; n_AP2γ^+/−^ juvenile_ = 6; n_WT adult_ = 5; n_AP2γ^+/−^ adult_ = 5; 3D neuronal reconstruction: n_WT juvenile_ = 4; n_AP2γ^+/−^ juvenile_ = 4; n_WT adult_ = 4; n_AP2γ^+/−^ adult_ = 5. [Two-way ANOVA and Repeated measures ANOVA, Statistical summary in Supplementary table 1] \*\*\**p*<0.001, ** *p*< 0.01; * *p*< 0.05. Abbreviations: WT, wild-type; AP2γ^+/−^, AP2γ heterozygous knockout mice.

**Supplementary figure 2:**
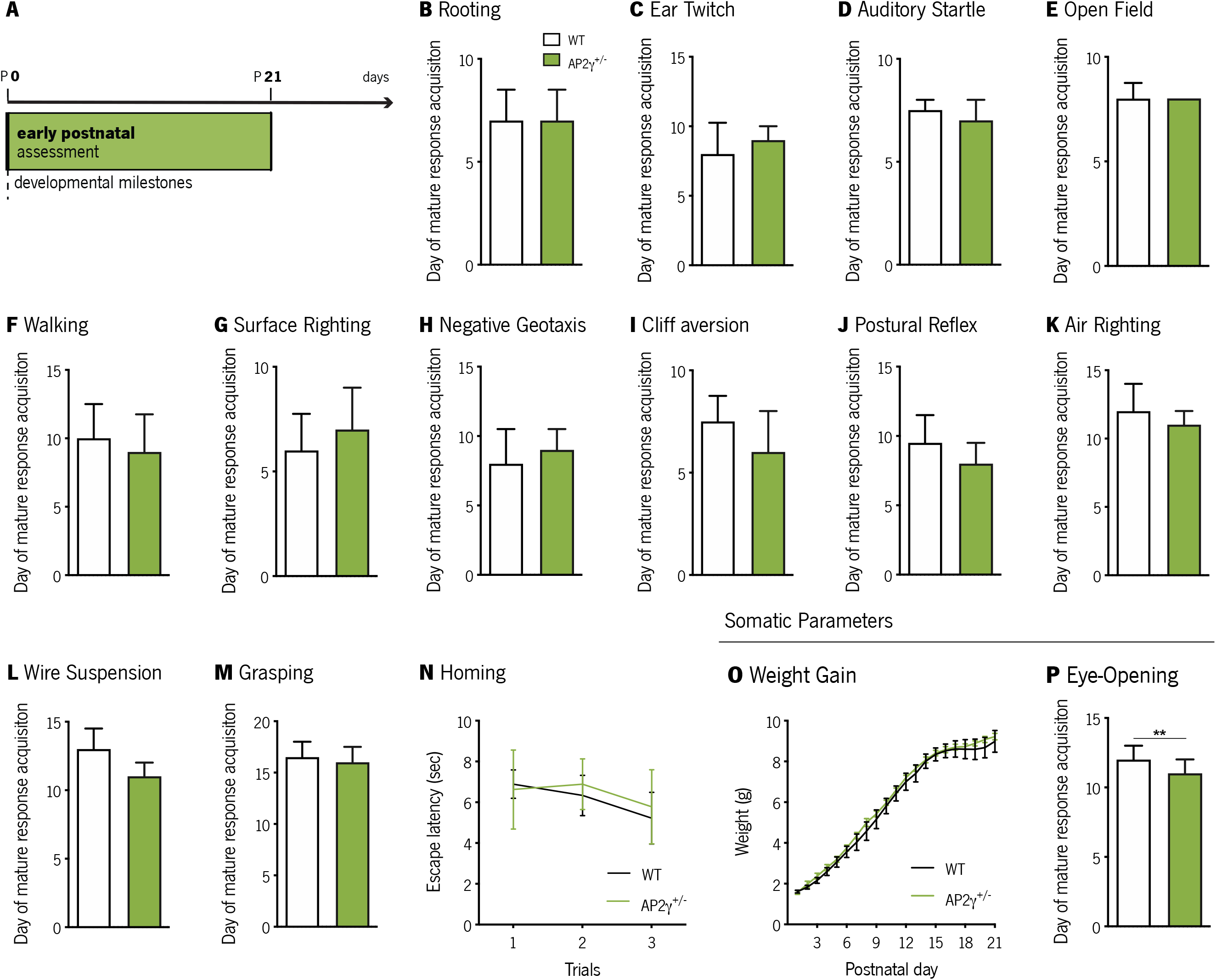
AP2γ constitutive and heterozygous deficiency does not impact on early postnatal development. (A) Timeline of early development assessment. (B-N) Set of established protocols to analyze the acquisition of mature responses, in neurobiological reflexes related to tactile (B and C) and auditory reflexes (D), motor function (E and F), vestibular system formation (G-K), strength (L and M), and olfactory maturation (N). Somatic parameters were also assessed. (O) Bodyweight gain from postnatal day (PND) 1 to PND 21 of WT and AP2γ^+/−^ mice. (P) Eye-opening day. For the weight gain pattern, data presented as mean SEM; for the remaining tests, data was plotted as median IQR. Sample Size: *n*_WT_ = 9; *n* AP2^+/−^ = 9. [Mann-Whitney and Repeated measures ANOVA, ** *p*< 0.01, Statistical summary in Supplementary table 1]. Abbreviations: WT, wild-type; AP2γ^+/−^, AP2γ heterozygous knockout mice; IQR, Interquartile range.

**Supplementary figure 3:**
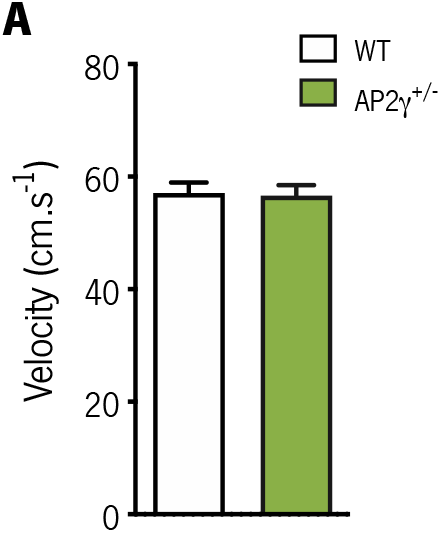
AP2γ deficiency does not impact on motor function in juvenile mice. (A) Average velocity assessed through the open field (OF) test in juvenile mice. Data presented as mean SEM. Sample Size: OF: *n*_WT_ = 13; *n*_AP2γ^+/−^_ = 11. [Student’s t-test, Statistical summary in Supplementary table 1]. Abbreviations: WT, wild-type; AP2γ^+/−^, AP2γ heterozygous knockout mice.

**Supplementary figure 4:**
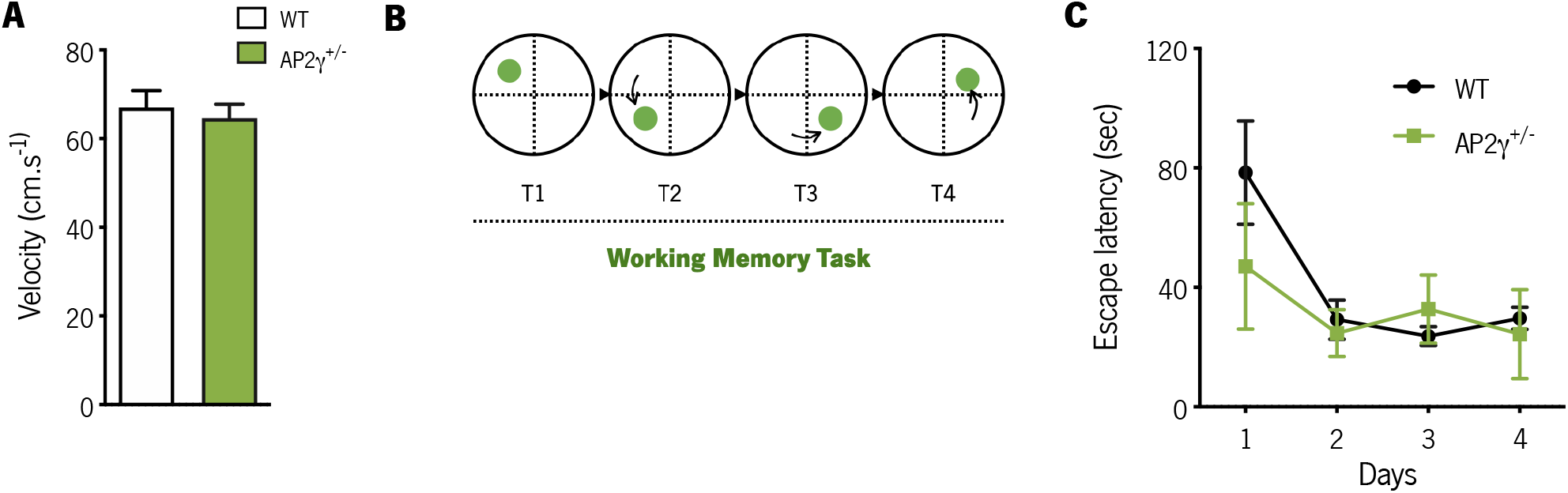
Deficiency in AP2γ transcription factor levels does not impact motor function, nor specific modalities of cognitive behavior. (A) Average velocity assessed through the open field (OF) test in adult mice. (B) Working memory task evaluated in the Morris water maze (MWM) test. Data presented as mean SEM. Sample size: OF: *n*_WT_ = 12; *n*_AP2γ^+/−^_ = 14; MWM: *n*_WT_ = 10; *n*_AP2γ^+/−^_ = 10. [Student’s t-test and Repeated measures ANOVA, Statistical summary in Supplementary table 1]. Abbreviations: WT, wild-type; AP2γ^+/−^, AP2γ heterozygous knockout mice.

**Supplementary figure 5:**
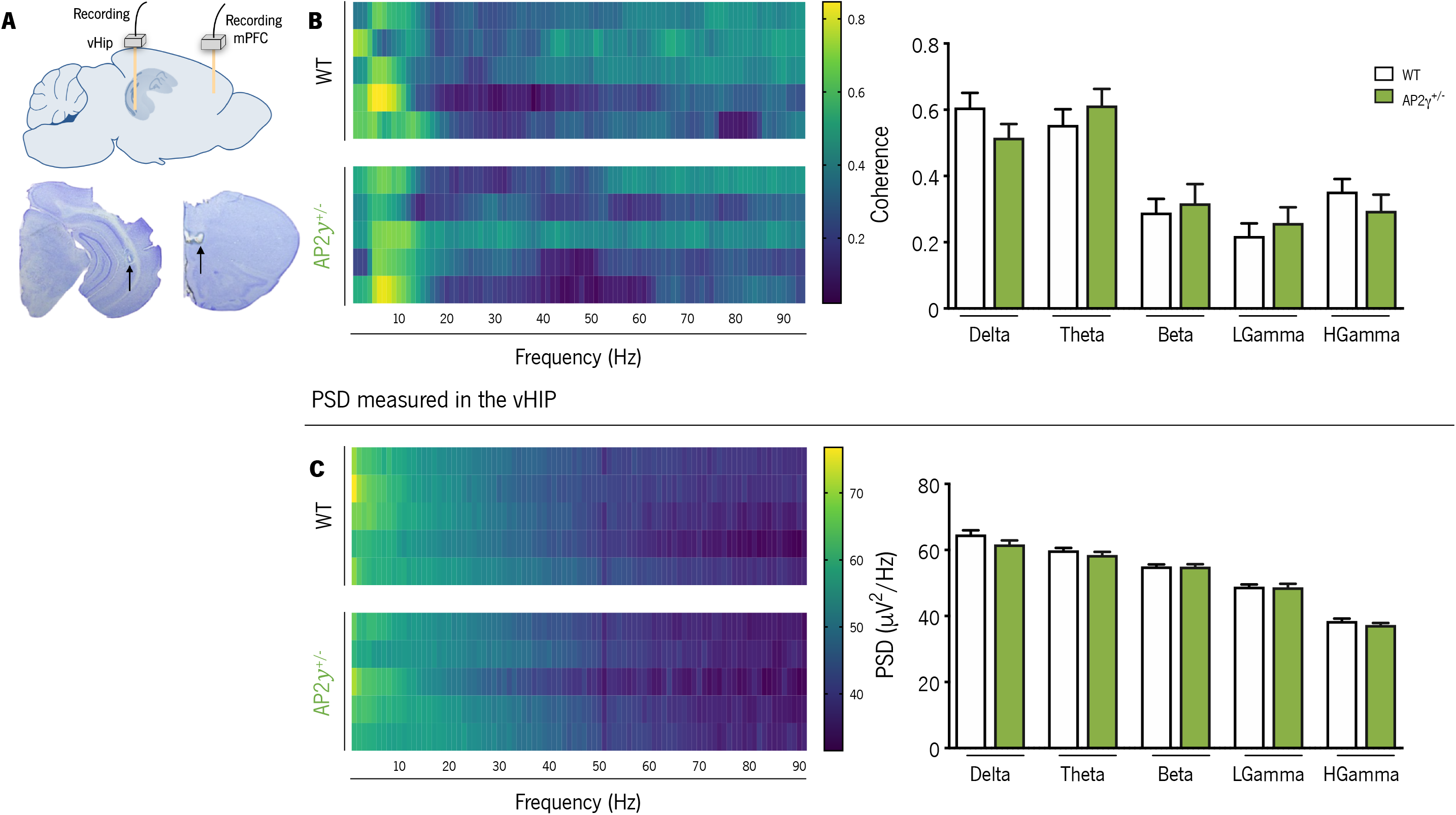
AP2γ deficiency does not impact neither on spectral coherence between the ventral hippocampus (vHip) and the medial prefrontal cortex (mPFC), nor the neuronal activity in each region. (A) Local field potentials (LFP) recording sites, with a depiction of the electrode positions (upper panel), and representative Cresyl violet-stained section (lower panel). (B) Spectral coherence (left panel) and group comparison for each frequency (right panel), between the vHip and mPFC of adult WT and AP2γ^+/−^ mice. (C) Power spectral density (PSD) (upper panel), and group comparison of the PSD values in the vHip for each frequency (lower panel). In spectrograms, each horizontal line in the Y-axis represents an individual mouse. Frequency bands range: delta (1-4Hz), theta (4-12 Hz), beta (12-20 Hz), low gamma (20-40 Hz), and High gamma (40-90 Hz). Data presented as mean SEM. *n*_WT_ = 5; *n*_AP2γ^+/−^_ = 5. [Student’s t-test, Statistical summary in Supplementary table 1]. Abbreviations: WT, wild-type; AP2γ^+/−^, AP2γ heterozygous knockout mice; vHip, ventral hippocampus; mPFC, medial prefrontal cortex.

